# Promoter-intrinsic and local chromatin features determine gene repression in lamina-associated domains

**DOI:** 10.1101/464081

**Authors:** Christ Leemans, Marloes van der Zwalm, Laura Brueckner, Federico Comoglio, Tom van Schaik, Ludo Pagie, Joris van Arensbergen, Bas van Steensel

## Abstract

It is largely unclear whether genes that are naturally embedded in lamina associated domains (LADs) are inactive due to their chromatin environment, or whether LADs are merely secondary to the lack of transcription. We show that hundreds of human promoters become active when moved from their native LAD position to a neutral context in the same cells, indicating that LADs form a repressive environment. Another set of promoters inside LADs is able to "escape" repression, although their transcription elongation is attenuated. By inserting reporters into thousands of genomic locations, we demonstrate that these escaper promoters are intrinsically less sensitive to LAD repression. This is not simply explained by promoter strength, but by the interplay between promoter sequence and local chromatin features that vary strongly across LADs. Enhancers also differ in their sensitivity to LAD chromatin. This work provides a general framework for the systematic understanding of gene regulation by repressive chromatin.

**Highlights:**

- Two promoter transplantation strategies elucidate the regulatory role of LAD chromatin
- LADs are generally repressive, but also highly heterogeneous
- LADs can impede both promoter activity and transcription elongation
- Promoters vary intrinsically in their sensitivity to LAD repression

## INTRODUCTION

Heterochromatin is generally defined as a compacted chromatin state, and is thought to repress the transcriptional activity of genes and mobile elements (Allshire and Madhani, 2017). For many individual genes in a variety of species this effect on transcription has been demonstrated. The evidence consists typically of two observations: (i) in its inactive state, the gene is naturally embedded in heterochromatin; (ii) the gene becomes active when a key component of the heterochromatin protein complex is removed. Despite many compelling anecdotal examples of genes that fit these criteria, genome-wide studies have rarely found that all genes within a certain heterochromatin type are activated when key heterochromatin proteins are removed or mutated. Rather, only a fraction of the heterochromatic genes become active after such perturbations (Bulut-Karslioglu et al., 2014; King et al., 2018; Pengelly et al., 2015; Penke et al., 2016; Santoni de Sio et al., 2012; Yokochi et al., 2009).

One possible explanation for this conundrum could lie in the redundancy of heterochromatin proteins, which may make it difficult to disable heterochromatin completely by depletion of a single protein. An alternative possibility is that many genes in heterochromatin are inactive due to other causes, for example due to the absence of an essential transcription factor (TFs) in the cell type that is studied. If this is true, then lifting the heterochromatic state would not be sufficient to release gene activity. It is generally not known which fraction of human genes in heterochromatin is actively repressed by the heterochromatic context, and which fraction is passively inactive due to lack of an essential activating signal.

Lamina Associated Domains (LADs), are among the most prominent heterochromatin domains in metazoan genomes (Gonzalez-Sandoval and Gasser, 2016; Luperchio et al., 2017; Van Steensel and Belmont, 2017). In mammalian cells, LADs are about 10 kb - 10 Mb in size and collectively cover about 30-40% of the genome (Guelen et al., 2008; Peric-Hupkes et al., 2010). LADs interact closely with the nuclear lamina (NL) and are enriched in the histone modifications H3K9me2 and H3K9me3, and in some cases H3K27me3. In mammals, several thousands of genes are located within LADs, and the majority of these genes is transcriptionally inactive. It is believed that LADs may form a repressive chromatin state. This notion is supported by experiments involving artificial tethering of genomic loci to the NL, which caused reduced expression of some, but not all, genes in the tethered loci (Finlan et al., 2008; Kumaran and Spector, 2008; Reddy et al., 2008). However, the regulation of mammalian genes *naturally* embedded in LADs is poorly understood. In particular, for very few of these genes it has been conclusively demonstrated that the lack of transcriptional activity is due to the heterochromatic state of LADs. This may be because critical protein components of LAD chromatin still have not been identified. Additionally, if the heterochromatic state of LADs indeed inhibits transcription, it may do so through several mechanisms: It may block initiation at the promoter, but also transcription elongation, and it may silence enhancers that are essential for promoter activity. Which of these mechanisms may play a role within LADs is not known.

Although the majority of genes in LADs is inactive, about 10% are expressed (Guelen et al., 2008; Wu and Yao, 2017). Analysis of such exceptions to the rule may provide valuable mechanistic insights. Possibly, these genes somehow are able to overrule the (putative) repressive LAD environment, for example if their promoters are extremely strong. Alternatively, LADs may be heterogeneous, and the local chromatin context of these active genes may be less repressive or even stimulatory. It is not known which of these mechanisms - which are not mutually exclusive - allow genes to be active inside LADs.

Here, we report a detailed analysis of gene repression mechanisms in LADs. In particular, we studied how promoter activity is controlled by LADs. For this purpose we made use of two orthogonal high-throughput genomic transplantation strategies. We systematically moved promoters from their native LAD location to a more neutral chromatin environment, and we also transplanted them to a wide range of locations including many LAD contexts. Combined with extensive computational analysis, the results reveal that indeed hundreds of promoters inside LADs are repressed by the local chromatin state. Moreover, the data demonstrate that features encoded in the promoter sequence, as well as strong variation in local LAD composition, determine the expression level of genes inside LADs.

## RESULTS

### Frequent repression of promoter activity in LADs

One approach to investigate whether a promoter embedded in a LAD in a certain cell type is repressed by the LAD chromatin context, is to insert the promoter into a reporter plasmid and transfect it into the same cells. This transfer to an episomal context may lead to increased activity of the promoter due to its release from the LAD environment.

We aimed to test hypothesis for many promoters in LADs. We recently reported Survey of Regulatory Elements (SuRE), a massively parallel reporter assay in which ~150 million random genomic DNA fragments are assayed for promoter activity in a transiently transfected plasmid (van Arensbergen et al., 2016). These genome-wide SuRE data include quantitative measurements of the activity of all annotated human promoters (**Supplementary Dataset S1**), thus offering an opportunity to systematically investigate whether LAD promoters are activated when moved into a plasmid reporter.

As a measure of promoter activity in the native chromatin context we used data obtained with the GRO-cap method (Core et al., 2014). We focused on 31,043 well-annotated promoters (see Methods) in human K562 cells, a widely studied leukemia cell line for which both SuRE and GRO-cap data are available. In inter-LAD regions (iLADs) promoter expression measured by the two methods show a substantial correlation (**Figure 1A**; Peason’s r = 0.70; Spearmans rank correlation rho = 0.76), in line with what was previously observed genome-wide (van Arensbergen et al., 2016). This correlation is weaker in LADs (**Figure 1B**; r = 0.53; rho = 0.52). More importantly, LAD promoters exhibit on average ~10-fold lower GRO-cap signals than iLAD promoters with matching SuRE activity (sliding window curves in **Figure 1B**). Furthermore, 58% of the 2,048 LAD promoters with SuRE activity (SuRE log_10_ expression values > 0) exhibited no detectable GRO-cap activity, compared to 24% of a set of 3,289 iLAD promoters that were selected to have a closely matching SuRE activity (**Supplementary Figure S1A, B**). Thus, promoters with the same intrinsic activity tend to be much less active when embedded in LADs compared to iLADs. This indicates that LADs are generally poorly conducive to promoter activity, compared to iLADs.

**Figure 1.**
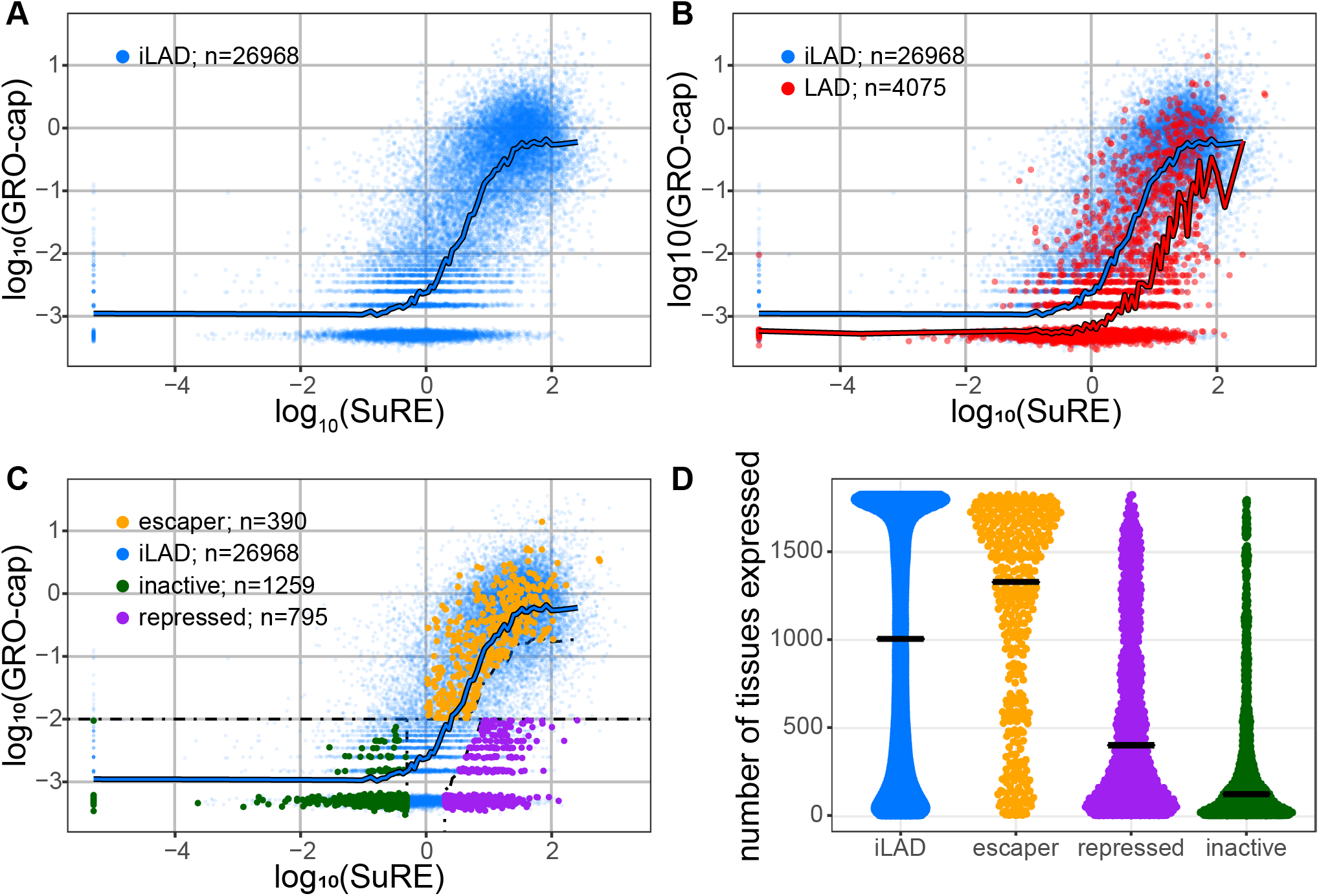
Promoters respond differently to their LAD environment. **(A)** Relation between promoter activity in the native context (GRO-cap) and in episomal plasmid context (SuRE) for all inter-LAD (iLAD) promoters (blue dots). Blue line shows sliding window average with a bin size of 501 promoters along the SuRE expression axis. GRO-cap data are from (Core et al., 2014); SuRE data are from (van Arensbergen et al., 2016). **(B)** Comparison of all promoters in LADs (red) and iLAD regions (blue). Red line shows sliding window average of LAD promoters, similar to blue line in (A). **(C)** Same as in (B), but with three classes of LAD promoters highlighted in green, purple and amber. Dotted lines depict cutoff lines used for the definitions (see Methods). **(D)** Promoter classes differ in their tissue expression distributions. Plot shows the number of tissues and cell types (out of 1829) in which promoter activity was detected by the FANTOM5 study ((DGT) et al., 2014). Each dot represents a promoter in either iLADs (blue) or one of the three LAD promoter classes (amber, purple, green).

### Three classes of promoters in LADs

Despite this clear general trend, not all LAD promoters respond similarly to their environment. Some exhibit high GRO-cap activity, suggesting that they can somehow escape the repressive influence of their LAD environment. To investigate this heterogeneity further, we defined three distinct categories of LAD promoters (**Figure 1C**): (i) *repressed* LAD promoters, which show >10-fold lower GRO-cap activity than typical iLAD promoters with similar SuRE activity; (ii) *escaper* LAD promoters, which exhibit GRO-cap activity that is similar to or even higher than that of iLAD promoters with matching SuRE activity; and (iii) *inactive* LAD promoters, which show very low SuRE and GRO-cap signals.

We initially interpreted these three classes of LAD promoters as follows. *Repressed* promoters have the potential to be active (i.e., all required transcriptional activators are present in the cell), but appear to be repressed by their native LAD context. In contrast, *escaper* promoters may carry features that render them less sensitive to the repressive effects of LADs, or they may reside in a LAD sub-region that is not repressive. Finally, *inactive* promoters appear to lack an essential activating signal, both in their native environment and in the plasmid context. This could be due to the absence of a critical activating transcription factor (TF) in the cell type studied. Alternatively, it could be because the promoter requires a distal enhancer that is not included in the SuRE reporter plasmid, and that is also not functional in the native LAD context.

Of the 2444 thus classified promoters in LADs, 33% are repressed,16% are escapers, and 52% are inactive. Interestingly, the three promoter classes show a difference in their overall tissue specificity (**Figure 1D**). Escaper LAD promoters are broadly expressed, and based on this many may be classified as promoters of housekeeping genes. In contrast, inactive LAD promoters are mostly highly tissue-specific, and repressed LAD promoters exhibit an intermediate pattern of tissue-specificity. These observations underscore that the three promoter classes are of distinct nature.

### Escaper promoters are locally detached from the NL

While escaper promoters are by definition located inside LADs, close inspection of the DamID data suggested that they are locally detached from the NL. On average, the DamID signals around these promoters are about 4-fold lower than for their upstream and downstream regions (**Figure 2A**). The detached region typically extends from about 10kb upstream to 10kb downstream of the TSS, although even beyond these distances the DamID signals remain somewhat lower than the average signal inside LADs. Virtually no such detachment was observed for the repressed and inactive promoter classes (**Figure 2A**). Of note, the ~20kb region around escaper promoters that appears to be detached from the NL is substantially larger than the size of the promoter itself or of the nucleosome-free region that is detected by DNase-seq (**Supplementary Figure S2A**).

**Figure 2.**
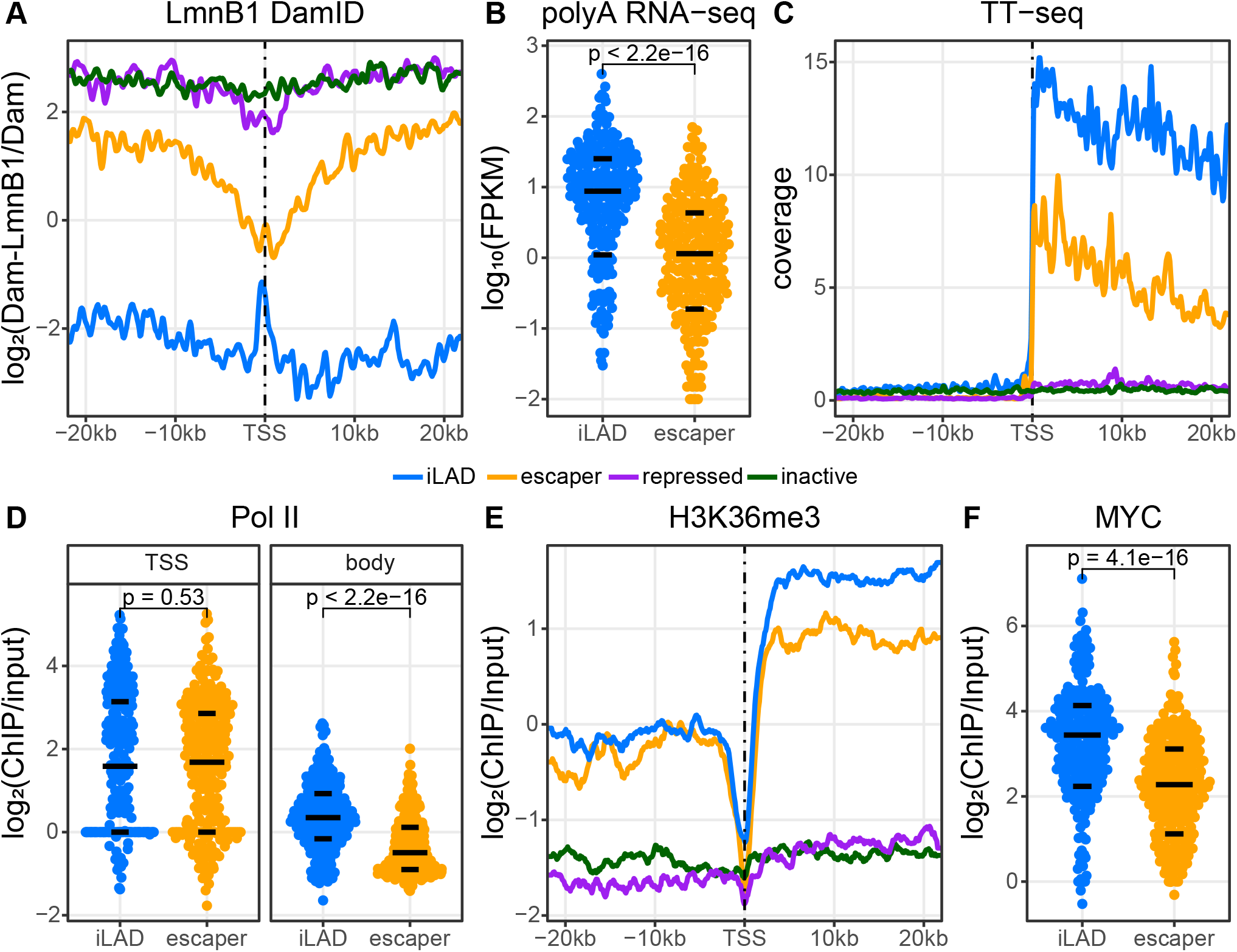
Properties of escaper promoters and their genes in native context. **(A)** Average NL interaction profile around each promoter class according to DamID-seq of Lamin B1. Note the local detachment of escaper promoters. Data are average of two independent experiments. Only genes were included that are at least 20 kb long, and for genes in iLADs, only those that do not have any other gene within 40 kb upstream of the TSS were used. **(B)** Total mRNA expression level of genes driven by escaper promoters compared to a set of genes driven by iLAD promoters with matching GRO-cap activity (blue). mRNA-seq data are from (Dunham et al., 2012). **(C)** Average TT-seq profile of each promoter set as in (B). Data from (Schwalb et al., 2016). **(D)** ChIP-seq signals of RNA Polymerase II around the TSSs (-50 to +300 bp; left panel) and along the gene bodies (+300bp from the TSS to +3000 bp downstream of the transcription termination site) of genes driven by each promoter set as in (B). Data are from (Dunham et al., 2012). **(E)** Average H3K36me3 signals along genes driven by each promoter set as in (B). Data are from (Schmidl et al., 2015). **(F)** ChIP-seq signals of c-Myc protein (Dunham et al., 2012) at each promoter set as in (B). Black horizontal lines in **B**, **D** and **E** depict 25th, 50th and 75th percentiles. P-values in B, D and E are from a Wilcoxon rank-sum test.

NL interactions are in part controlled by the histone modifications H3K9me2 and H3K9me3 (Bian et al., 2013; Harr et al., 2015; Kind et al., 2013; Towbin et al., 2012). Analysis of ChIP-seq data shows that escaper promoters indeed show a local decrease of these marks (**Supplementary Figure S2B, C**). However, the other promoter classes show a nearly similar local decrease of H3K9me2 and H3K9me3, suggesting that the detachment of escaper promoters from the NL cannot be solely explained by the absence of these marks. The local detachment from the NL might facilitate the transcriptional activity of escaper promoters, although we cannot rule out that it is a secondary consequence of their activity (Chuang et al., 2006; Therizols et al., 2014).

### Partially impaired transcription elongation from escaper promoters in LADs

Since escaper promoters are active, we expected that the corresponding genes would produce similar amounts of mRNA as their iLAD counterparts. Surprisingly, we found that this is not the case. Escaper genes produce on average 5-fold less mRNA than a set of iLAD genes with the same promoter activity distribution as measured by GRO-cap (**Figure 2B**). Escaper genes also produced less mRNA than a set of iLAD genes matched for promoter activity as measured by SuRE (**Supplementary Figure S2D**).

To investigate this discrepancy between promoter activity and mRNA yield, we examined marks of transcription elongation. The amount of nascent RNA as detected by the TT-seq method (Schwalb et al., 2016) was lower along the gene bodies of escaper genes (**Figure 2C**), as was the occupancy of RNA Polymerase II (Pol II) (**Figure 2D**). Also H3K36me3, another mark of elongation (Wagner and Carpenter, 2012), was lower along escaper gene transcription units (**Figure 2E**). In each of these analyses we compared the escaper genes to a set of iLAD genes with matched promoter activities according to GRO-cap. We note that these elongation marks are reduced along the entire length of the genes; while there is some indication of progressive loss of TT-seq signals towards the 3’ ends, this is only marginally more pronounced than in matching iLAD genes (**Figure 2C**). Together, these results point to inefficient transition of Pol II from initiation to elongation, rather than to random abortion of elongation along the gene body.

In a survey of publicly available ChIP-seq data for proteins that might explain the attenuated elongation of escaper genes, we found that Myc binding is nearly 2-fold weaker at escaper promoters compared to activity-matched iLAD promoters (**Figure 2F**). Myc has previously been reported to stimulate the release of paused polymerase from promoters (Rahl and Young, 2014). The reduced binding of Myc may in part explain the relatively poor elongation of escaper genes, but it is also possible that association with the NL (or a chromatin feature linked to this association) impedes the initiation-to-elongation transition.

### Testing repressed and escaper promoters in many chromatin contexts

One possibility is that the different activity levels of repressed and escaper promoters inside LADs is encoded in their proximal sequence. If this is true, repressed promoters should remain inactive when transplanted to a different LAD, and escaper promoters should remain active when moved to a different LAD. Alternatively, the differences between repressed and escaper promoters may reflect local differences in LAD context, with some sub-LAD regions being more repressive than others. If this is the case, then the activity of both repressed and escaper promoters should strongly depend on the precise LAD context in which they are located.

To discriminate between these two hypotheses, we inserted several escaper and repressed promoters into a large number of different genomic locations (both LAD and iLAD) by means of the Thousands of Reporters Integrated in Parallel (TRIP) method (Akhtar et al., 2013). Specifically, we cloned representative promoters of each class into a common reporter construct, and generated K562 cell pools in which the reporters were integrated randomly into hundreds of genomic locations, including many LADs. We mapped the integration sites by inverse-PCR and high-throughput sequencing (Akhtar et al., 2014). Because we marked each copy of the reporter with a unique random barcode in its transcription unit, we could determine the expression level of all integrations in parallel by high-throughput sequencing and counting of each barcode in mRNA from the cell pools. This strategy enabled us to study the effect of many different LAD and iLAD contexts on individual promoters.

We performed these TRIP experiments with three repressed promoters and three escaper promoters. In addition, we included the promoter of the *PGK* gene, a housekeeping gene that is located in an iLAD (**Supplementary Table S1**). For these seven promoters combined, we obtained 12,812 integrations (629-3,514 per promoter) that could be mapped to unique genomic locations (**Supplementary Figure S3**). Of these, 23% were located inside a LAD. This is less than may be expected by random chance, because LADs occupy 43% of the genome in K562 cells. This may reflect an integration bias of the PiggyBac vector (de Jong et al., 2014), but also a relatively poor mappability of integrations inside LADs due to a higher repeat content. Nevertheless, we obtained 123-737 integrations in LADs per promoter, providing ample statistical power, as will be shown below. Expression of the barcoded reporter transcripts in the cell pools carrying these integrations were analyzed in two independent biological replicates each.

### Repressed and escaper promoters differ in sensitivity to LAD context

All six promoters originating from LADs showed strong variation in expression levels, depending on their integration site (discussed in more detail below). For the three promoters of the repressed class, the median expression level was about 43-130 fold lower within LADs than within iLADs (**Figure 3A-C**), underscoring the repressive potential of LADs. For the escaper promoters, however, the difference between LAD and iLAD integrations was much less pronounced (4-20 fold; **Figure 3D-F**). Indeed, a linear regression model of reporter expression as function of Lamin B1 DamID signal and promoter class (escaper or repressed) showed a significant interaction between promoter class and Lamin B1 DamID levels (**Figure 3H**), indicating that repressed promoters and escaper promoters respond differently to the LAD environment.

**Figure 3.**
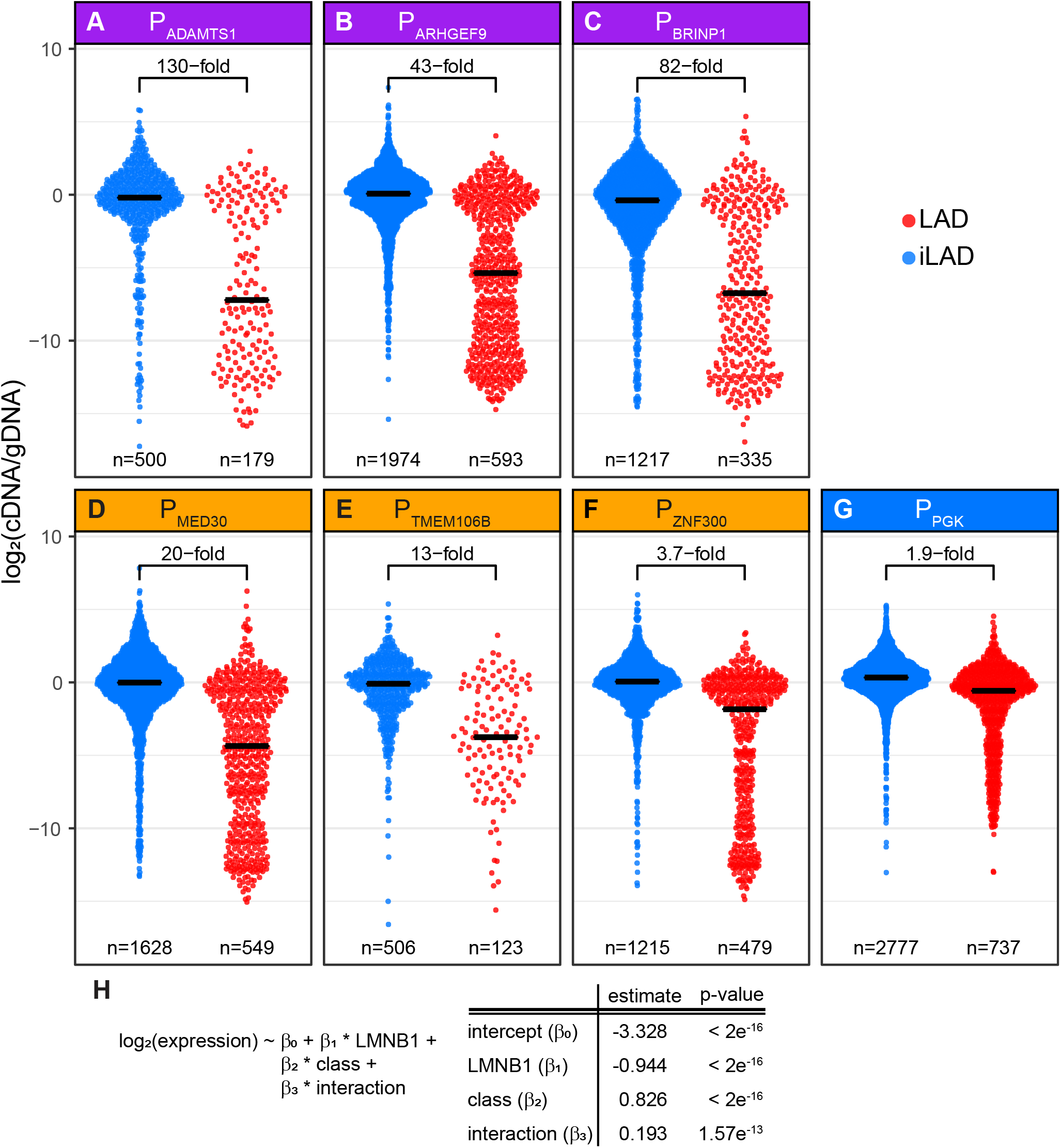
Effects of LAD context on expression of integrated reporters driven by various promoters. **(A-G)** Distributions of expression levels of barcoded reporters integrated in iLADs (blue) and LADs (red). Reporters were driven by the indicated LAD promoters from the repressed (purple, A-C) or escaper (amber, D-F) class, or by the iLAD-derived PGK promoter (blue, G). Horizontal black bars indicate medians; the fold-difference between the medians in iLADs and LADs are indicated. n denotes the number of integrated reporters assayed. Data are averages of two independent experiments. **(H)** Summary of a linear regression model of the expression levels of the integrated reporters as function of the local Lamin B1 DamID signal, the promoter class (escaper or repressed), and the interaction between promoter class and local DamID signal. All three terms contribute significantly to the model. Only LAD integrations from panels A-F were used in the model.

Surprisingly, the *PGK* promoter showed only a 2-fold difference in median expression between LAD and iLAD contexts (**Figure 3G**), indicating that it is largely refractory to LAD environments. This underscores that promoters differ in their sensitivity to LAD contexts, and suggests that the *PGK* promoter has strong escaper-like properties.

### Promoter strength does not explain differential sensitivity to LAD context

We considered the possibility that escaper promoters are intrinsically stronger than repressed promoters, and thereby may overrule the repressive environment of LADs more effectively. To test this hypothesis, we measured the activity of the promoters in episomal plasmid context, i.e., not embedded in any genomic chromatin environment. We considered this to be the most neutral environment that could reasonably be obtained. For the most precise estimate of promoter activity, we constructed mini-libraries of ~100 barcoded reporter plasmids for each promoter, mixed these in equal amounts, and transienty transfected the entire mix into K562 cells. Barcodes were then counted in mRNA, and normalized to barcode counts in the plasmid DNA pool. The representation of each promoter by ~100 random barcodes ensures average expression levels that are not biased by barcode sequence, and that are minimally affected by random noise, providing an accurate estimate of the relative activities of the promoters.

The results of these measurements show that the three escaper promoters are not systematically stronger than the repressed promoters (**Figure 4**). Thus, the intrinsic promoter strength does not account for the different responses of repressed and escaper promoter to LAD contexts. However, the *PGK* promoter is the by far the strongest of the seven promoters; possibly this contributes to its insensitivity to LAD environments.

**Figure 4.**
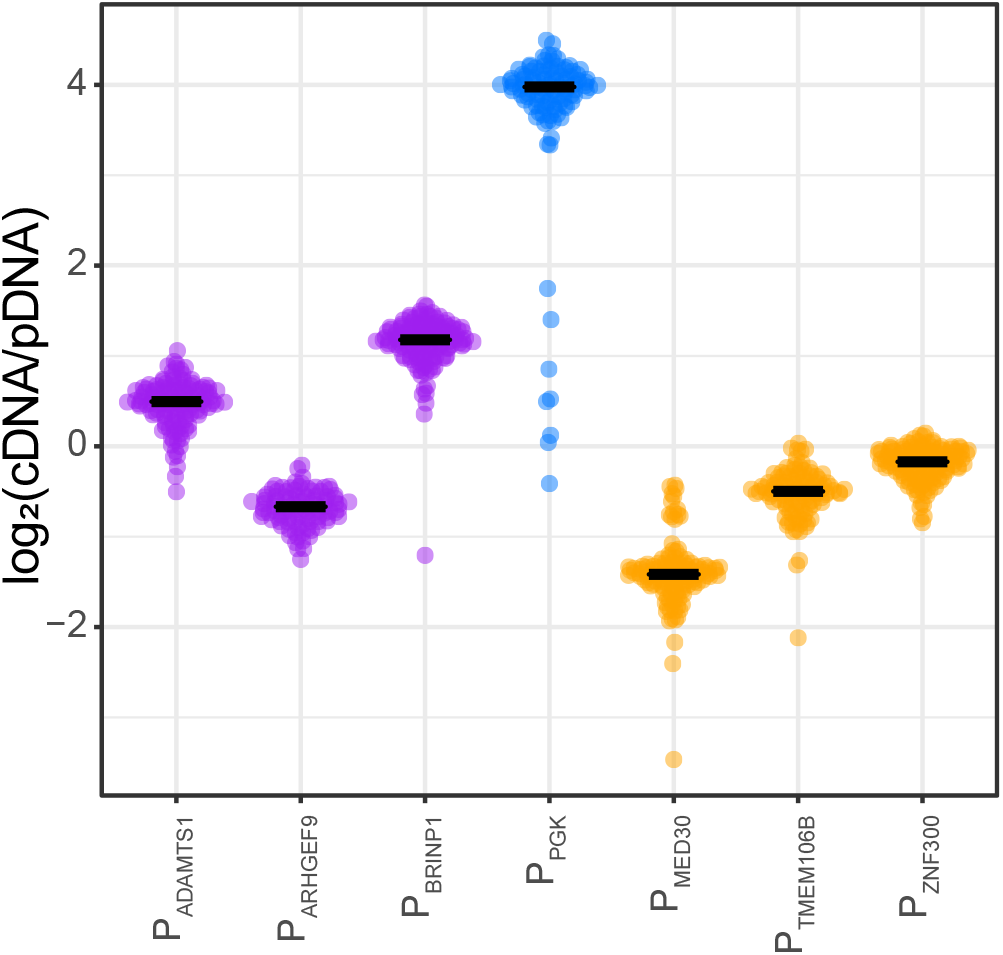
Precise measurement of promoter activities in episomal plasmid context. Each indicated promoter was cloned in the same promoter-less reporter plasmid with about 100 different random barcodes. A pool of the resulting ~700 plasmids was transiently transfected into K562 cells. Barcodes were counted in cDNA and plasmid DNA isolated from these cells after 2 days. The plot shows the distribution of expression levels of all barcodes sorted by promoter; horizontal lines depict medians. Data are average of two independent experiments.

We note that these transient transfection experiments also provide an estimate of the variance in expression levels that may be caused by differences in barcode sequences and other sources of noise. In the transient transfections we observed unimodal distributions with SD = 0.35 ± 0.25 (n = 7 promoters). This contrasts starkly with the multi-modal and much broader distributions (SD = 3.66 ± 0.87; n = 7) obtained with integrated reporters (**Figure 3A-G**). Thus, most of the variance observed with integrated reporters is not explained by barcode differences or technical noise, but rather by chromatin context.

### Chromatin features that correlate with promoter activity within LADs

Aside from the overall differences between LADs and iLADs, there is substantial variation in the TRIP reporter expression levels *within* LADs. This implies that LADs are heterogeneous structures, and that local chromatin features within LADs can substantially affect reporter activity. We note that within LADs the three repressed promoters showed a broader distribution of reporter expression levels than the three escaper promoters (**Figure A-F3**). Thus, compared to escaper promoters, repressed promoters are not only more sensitive to global LAD/iLAD differences, but also to local chromatin differences within LADs.

In order to investigate which features may explain the variation within LADs, we took a statistical learning approach. We compared our maps of reporter activity to a collection of available high-quality epigenomic maps from K562 cells (**Supplementary Tables S2, S3**). We considered both the signal strength of these epigenome features at the integration sites, and the distance to the nearest peaks or domains of these features. Note that these features were mapped in the absence of the integrations and can therefore not be the consequence of the integrations; conversely, it was previously shown that integrated reporters generally adopt the local chromatin state (Corrales et al., 2017), although exceptions cannot be ruled out. To identify the most likely candidate chromatin features to explain our reporter activities, we applied a feature selection algorithm that combines a lasso linear regression model with bootstrapping. This approach identified a set of features that statistically explain the reporter expression levels, with an estimate of their relative contribution to the predictive power of the model.

We restricted this analysis to integrations inside LADs. Collectively, chromatin features could explain nearly half of the variance in reporter expression for repressed promoters (R^2^ = 0.49 ± 0.05, mean ± SD across 1,000 samplings) and significantly less for escaper promoters (R^2^ = 0.35 ± 0.05; P = 8.16e-31, two-sided Wilcoxon test of R^2^ values) (**Figure 5A**). To corroborate these findings, we fitted promoter-specific models on all integrations (both LAD and iLAD). R^2^ values of promoter-specific model shows that results in **Figure 5A** are not driven by a single promoter, but this observation can be contributed to each of the 3 representitive promoters in each class (**Supplementary Figure S4**). These results underscore our previous conclusion that escaper promoters are less sensitive to their chromatin environment than repressed promoters.

**Figure 5.**
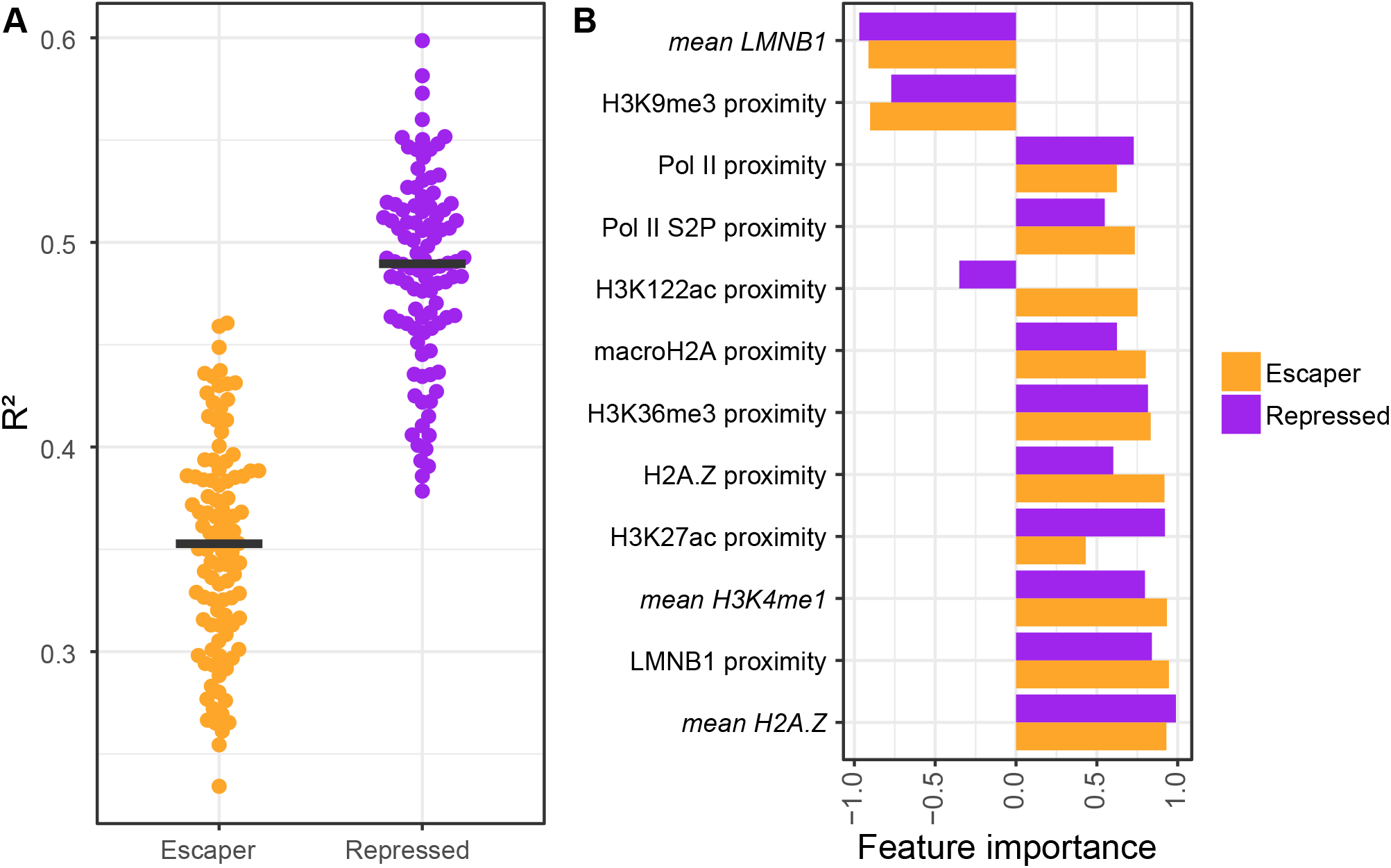
Modeling TRIP expression levels as function of LAD chromatin features. **(A)** R^2^ values of 100 lasso regression models of TRIP data for escaper and repressor promoters (see Methods). Each dot represents a model. Black lines represent the median R^2^ values. **(B)** Feature importance analysis of the most predictive chromatin features. Feature importance (x-axis) represents the fraction of bootstrap-lasso models in which a feature contributed significantly to the model performance. Negative and positive values of feature importance reflect negative and positive coefficient values, respectively. Mean signal intensities of chromatin features in a window around the integration site (5kb for ChIP, 10kb for DamID) as well as the proximity to the nearest peak of the same features were taken into account. The total list of features that was tested is listed in **Supplementary Table S2**. Only data from integrations inside LADs of the three repressed and three escaper promoters were used.

The chromatin features that were most predictive of reporter expression in LADs were generally shared between the two promoter classes, although we observed some quantitative differences in predictive power of individual features (**Figure 5B**). The LMNB1-DamID signal at the site of integration was one of the strongest predictors and negatively correlated with reporter activity. This signal was previously shown to be tightly linked to NL contact frequency (Kind et al., 2015), suggesting that reporters are more potently repressed when they are inserted in regions that are more stably associated with the NL. The local level of the histone variant H2A.Z is among the strongest positive predictors of reporter activity for both promoter classes. Mammalian H2A.Z generally marks active enhancers and promoters and is thought to promote binding of TFs and cofactors by destabilizing nucleosomes (Hu et al., 2013; Ku et al., 2012; Li et al., 2012). Also noteworthy is the predicted opposite effect of nearby H3K122ac peaks on escapers and repressed promoter classes. This this histone mark has been linked to a distinct type of enhancer that typically lacks H3K27ac (Pradeepa et al., 2016). Possibly this type of enhancer preferentially activates escaper promoters.

Together, these results underscore that escaper promoters are generally less responsive to chromatin heterogeneity inside LADs, and the modeling identifies the most likely chromatin features that may affect the activity of overlapping or nearby promoters.

### Promoter repression in H3K27me3 domains

We wondered whether the differential sensitivity of the seven promoters is specific for LADs, or also applies to other types of heterochromatin. We focused on domains of H3K27me3, which represent heterochromatin associated with Polycomb repressive complexes. In K562 cells 80% of the H3K27me3 domains are located outside of LADs. Interestingly, within these regions the integrated promoters also show striking quantitative differences in repression (**Figure 6A-G**). Generally, these differences correlate with those observed in LADs, but the degree of repression is systematically about 2-5 fold less in H3K27me3 domains compared to LADs (**Figure 6H**). Like in LADs, in H3K27me3 domains the three escaper promoters are less repressed than the three promoters of the LAD-repressed category, and the PGK promoter is essentially insensitive to H3K27me3 chromatin. In particular the ADAMTS1 and BRINP1 promoters show a broad range of expression levels inside H3K27me3 domains, pointing to a strong heterogeneity of these domains.

**Figure 6.**
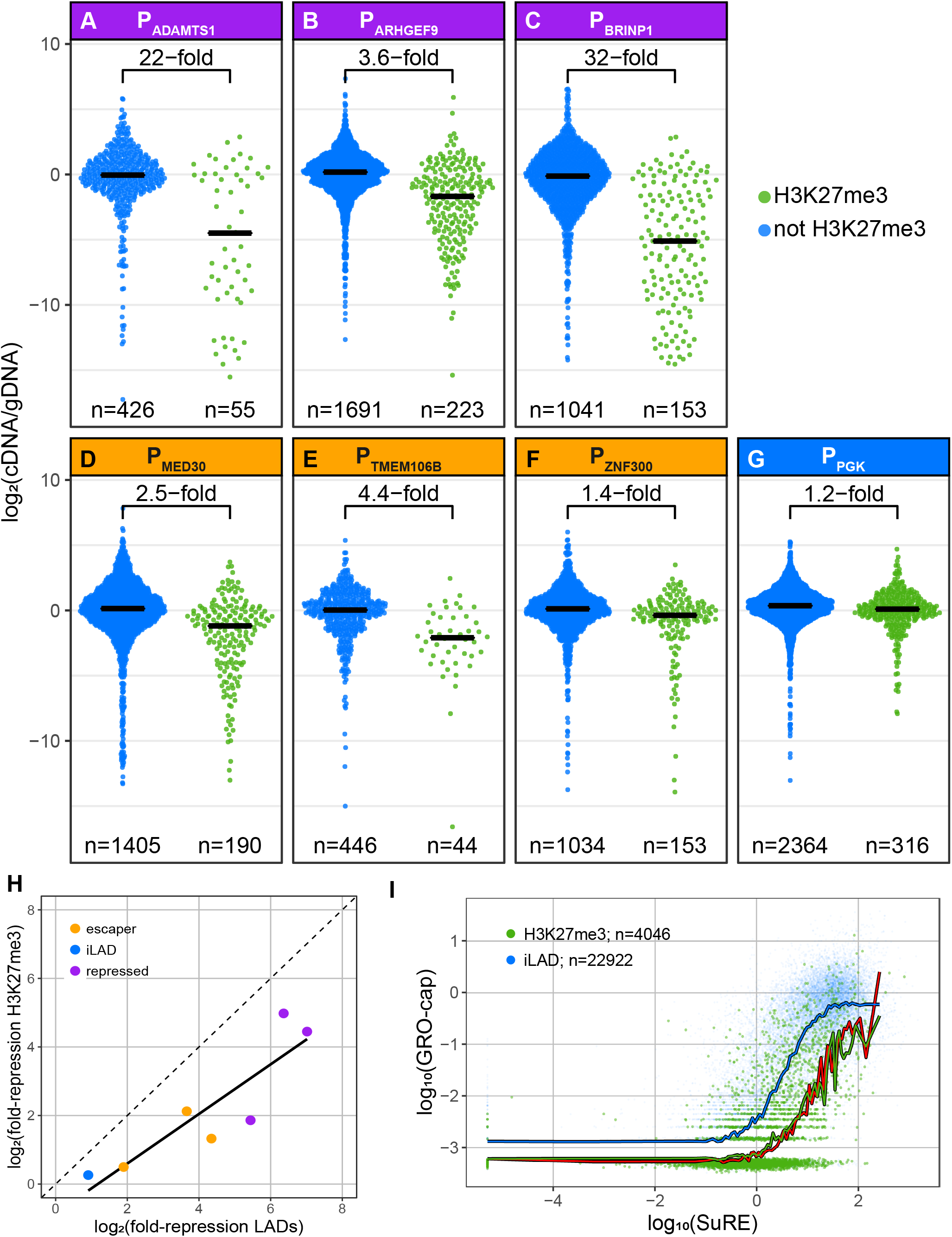
Effects of H3K27me3 domain context on promoter activity. **(A-G)** TRIP analysis of the same promoters as Figure 3 A-G, for reporters integrated in iLAD regions marked by H3K27me3 (green) or not(blue). **(H)** Correlation between repressive effect of LADs (fold difference between LAD and iLAD median expression, Figure 3A-G) and H3K27me3 domains (fold difference between H3K27me3 domain and non-H3K27me3 domain median expression, Figure 5A-G) for the seven promoters tested by TRIP. **(I)** SuRE versus GRO-cap analysis as in Figure 1A-B, for iLAD promoters located in H3K27me3 domains (green) compared to iLAD promoters not located in H3K27me3 domains (blue). Red curve shows LAD promoter trend line as in Figure 1B.

To complement these TRIP results, we revisited the SuRE and GRO-cap data and investigated the activity of iLAD promoters that are naturally located in H3K27me3 chromatin (**Figure 6I**). On average, promoters in H3K27me3 domains show a roughly 10-fold lower GRO-cap activity than SuRE-matched promoters located outside H3K27me3 domains (and outside LADs). This is similar to what we observed in LADs (see Discussion).

### Effects of LADs on enhancer activity

Finally, we explored the effects of LAD context on enhancers. We previously reported that SuRE can also detect enhancer activity, by virtue of the fact that enhancers typically act as transcription start sites and produce RNA in proportion to their activity (van Arensbergen et al., 2016). Similarly, in the native context, enhancers generate transcripts that can be detected by GRO-cap (Core et al., 2014). Hence, to investigate whether LADs might affect enhancer activity, we repeated the SuRE versus GRO-cap analysis, but now for enhancers (**Figure 7A**; **Supplementary Dataset S3**). This showed that LAD enhancers exhibit on average ~3-fold lower GRO-cap signals than iLAD enhancers with matching SuRE activity. Thus, like promoters, enhancers inside LADs appear generally repressed. It is possible that we underestimate the average magnitude of repression, because for enhancers the SuRE and GRO-cap signals are much closer to background levels than for promoters.

**Figure 7.**
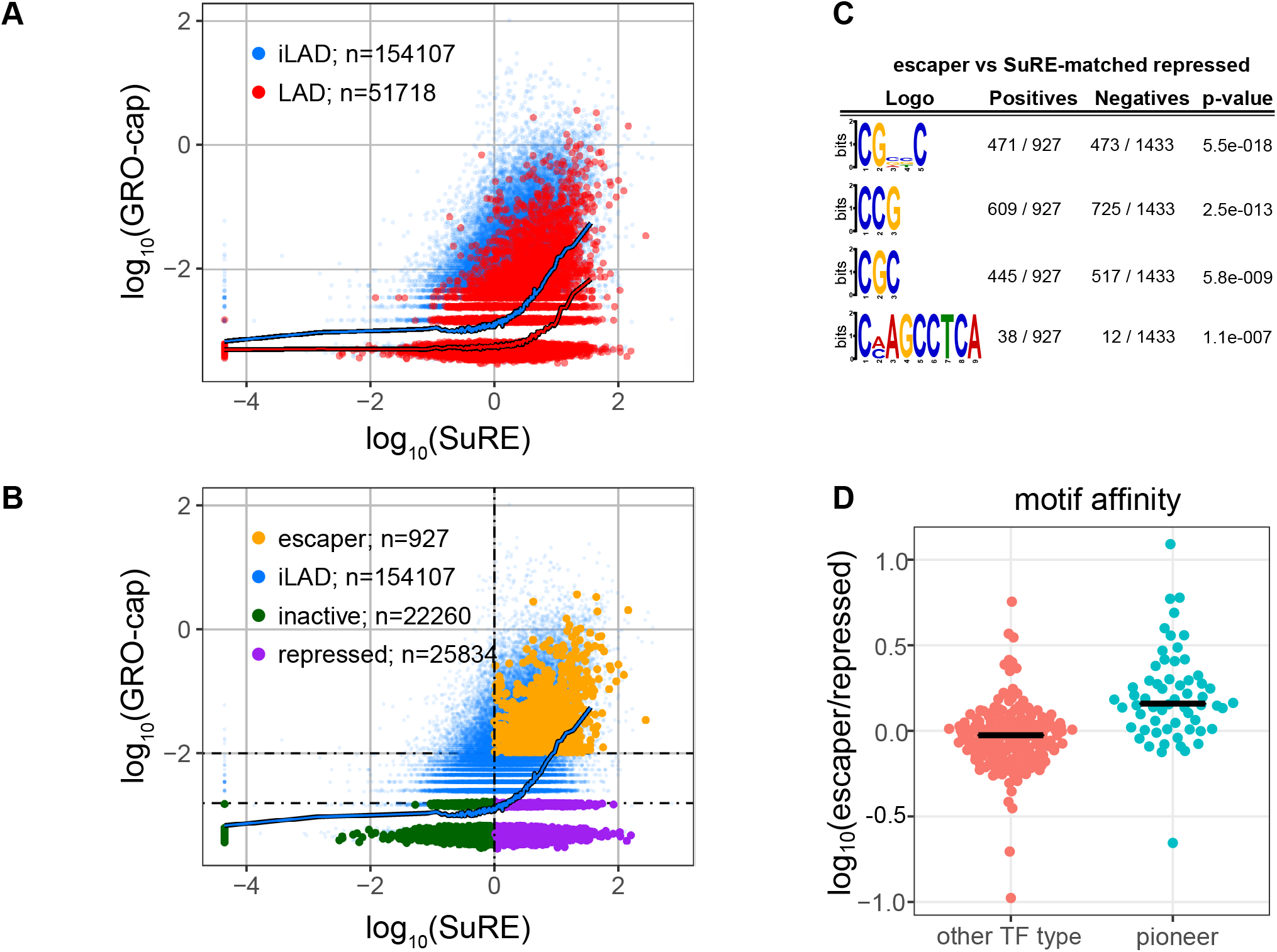
LAD repression of enhancer RNA production; effect of pioneer TF binding sites. **(A)** Comparison of enhancers in LADs (red) and iLADs (blue), similar to Figure 1B. Each dot represents a previously annotated enhancer (Ehsani et al., 2016). **(B)** Same as in (A), with three classes of LAD enhancers highlighted in green, purple and amber. Dotted lines depict cutoff lines used for the definitions (see Methods). **(C)** Sequence logos of enriched motifs in escaper enhancers compared to a set of repressed enhancers matched on SuRE activity. Motifs were identified by *DREME* (Bailey, 2011). **(D)** Fold-difference in median predicted affinity of enhancers for TFs. Each dot represents one TF that is detectably expressed in K562, separated into pioneer and non-pioneer TFs according to (Ehsani et al., 2016). Predicted affinity of each TF for each enhancer was calculated using the *AffinityProfile* from the *REDUCE Suite* (Roven and Bussemaker, 2003).

### Sequence motifs linked to enhancer sensitivity to LAD repression

Similar to escaper promoters, a subset of enhancers in LADs shows high GRO-cap signals that are more typical of enhancers in iLADs. Furthermore, some enhancers with very low GRO-cap signals were highly active in SuRE, while others were virtually inactive in SuRE. We therefore took a similar approach as for promoters and defined three classes of enhancers: inactive, repressed and escaper enhancers (**Figure 7B**). The large numbers of enhancers in each class provided sufficient statistical power to search for sequence motifs that may explain the difference in activity of escaper and repressed enhancers (a similar analysis on the much lower numbers of promoters lacked statistical power). *De novo* motif analysis comparing these two enhancer classes, corrected for SuRE activity, identified three prominent motifs enriched in escaper enhancers that all carry CpG dinucleotides, and one infrequent motif of unknown nature (**Figure 7C**). The CpG dinucleotide enrichment cannot be attributed to CpG islands, because very few of the escaper and repressed enhancers overlap with CpG islands (22 out of 927 and 15 out of 1433, respectively). No motifs were found that could be linked to specific TFs.

These results suggest that no single TF can account for the different responses of escaper and repressed enhancers to LADs. Instead, a broad class of TFs may contribute collectively to this difference. Prominent candidates are pioneer TFs, which are able to activate promoters inside condensed chromatin (Zaret and Carroll, 2011). We therefore tested whether cognate motifs for known and predicted pioneer TFs (Zaret and Carroll, 2011) collectively are enriched in escaper enhancers compared to repressed enhancers. Indeed, we observed a modest but statistically significant enrichment, suggesting that pioneer TFs collectively help escaper enhancers to overcome the repressive LAD environment (**Figure 7D**).

## DISCUSSION

In the past two decades, a wide diversity of genome-wide mapping methods (Dirks et al., 2016; Kelsey et al., 2017; Rivera and Ren, 2013; Zentner and Henikoff, 2014) has yielded a wealth of descriptive epigenome maps. Analysis of these data has uncovered extensive correlations between gene activity and many chromatin features. A current challenge is to move from these genome-wide correlations to a detailed understanding of causal relationships, while maintaining a genome-wide perspective. This requires systematic perturbation approaches (Catarino and Stark, 2018; Stricker et al., 2017). Here, we developed a general framework, centered around two genome-wide perturbation methods, to dissect causal relationships between local chromatin state and gene activity. The combined application of SuRE and TRIP provides detailed insights into the interplay between promoters and the local chromatin environment, here illustrated for LADs.

Previous TRIP experiments with two promoters (one from an iLAD, and one synthetic) had suggested that LADs form a repressive environment (Akhtar et al., 2013). Furthermore, it has been shown that some genes can be downregulated by artificial tethering to the NL (Dialynas et al., 2010; Finlan et al., 2008; Kumaran and Spector, 2008; Reddy et al., 2008). It remained unclear, however, whether genes that are naturally located in LADs are also repressed. Here, we identified hundreds of promoters that are repressed in their native context inside LADs, as evidenced by their activation upon transfer to an episomal plasmid in the same cells. Furthermore, for three promoters from the repressed class we confirm by TRIP that they indeed are extremely sensitive to a typical LAD environment, leading to ~40-130 fold reduction in their median expression levels compared to iLAD contexts.

Escapers, i.e., promoters that are active inside LADs have been observed before (Guelen et al., 2008; Wu and Yao, 2017), but it was not known how they overcome the repressive LAD environment. Our results identify two complementary determinants. First, we find that escaper promoters are intrinsically more resistant to the repressive LAD context than repressed promoters. This resistance is not generally due to a higher intrinsic transcriptional activity, although in some instances this may contribute. Instead, we suggest that escaper promoters contain sequence motifs that recruit a class of TFs that are somehow more effective in overcoming LAD repression. In addition, our analysis of escaper enhancers revealed a general enrichment of motifs that bind known and predicted pioneer TFs. It is possible that escaper promoters also rely on pioneer TFs, but we may have lacked statistical power to detect any enrichment of pioneer TF motifs in escaper promoters. Future experimental studies may investigate the contribution of pioneer TFs more directly. This may be challenging, however, because the escaper effect most likely does not depend on individual TFs, but rather on the combined effect of multiple TFs.

Our TRIP data also show that the activity of escaper promoters is higher in LAD regions where NL interactions as detected by DamID are weaker and when various active marks are nearby. Escaper promoters were previously found to carry such features (Wu and Yao, 2017), and our data are in agreement with this. DamID signal strength is known to correlate with NL contact frequencies (Kind et al., 2015), suggesting that escaper promoters may be active only when not contacting the NL. Importantly, our TRIP experiments help to interpret such correlative data in terms of causality, because these experiments show that the activity of escaper promoters is higher when they are inserted in or near regions that carried these features *prior to* integration of the reporters. It is thus highly likely that the activity of escapers in the native context is also at least partially facilitated by these features.

About half of all promoters located in LADs cannot be activated by transfer to an episomal plasmid. The lack of activity of these promoters is most likely explained by the absence of an activator. This missing activator may be either a promoter-binding TF that is not expressed in K562 cells, or a distal enhancer that is inactive or unable to contact the promoter. The repressive environment of LADs may provide an additional ‘lock’ to keep these genes inactive (Peric-Hupkes et al., 2010). However, it is also possible that some inactive genes associate passively with the NL as a consequence of their lack of activity. In support of this, it has been shown that forced activation of genes in LADs can lead to their detachment from the NL (Therizols et al., 2014).

Finally, our analysis of SuRE and GRO-cap data indicate that H3K27me3 domains are generally equally repressive domains as LADs. Our TRIP data confirm the repressive effect of H3K27me3 but surprisingly, the promoters tested by TRIP appear less sensitive to H3K27me3 contexts than to LAD contexts. One interesting possibility is that promoters naturally embedded in H3K27me3 domains have evolved to be particularly sensitive to this chromatin environment. We cannot rule out, however, that the PiggyBac transposon sequences used in TRIP partially counteract the repressive effect of H3K27me3 (more than the repressive effect of LAD chromatin), for example by specifically impeding the spreading of H3K27me3 chromatin into the promoters. Future studies may unravel the molecular architecture of LADs and the molecular mechanisms of gene repression and escape inside these domains.

## ACKNOWLEDGMENTS

We thank the NKI Genomics Core Facility for technical support and members of the Division of Gene Regulation for helpful discussions. Supported by National Institutes of Health Common Fund 4D Nucleome Program (Grant U54DK107965), ERC Advanced Grants 293662 and 694466, and ZonMW TOP to BvS; and an Early Postdoc mobility fellowship from the Swiss National Science Foundation to FC. The Oncode Institute is supported by KWF Dutch Cancer Society.

## SUPPLEMENTARY FIGURE LEGENDS

**Supplementary figure S1.**
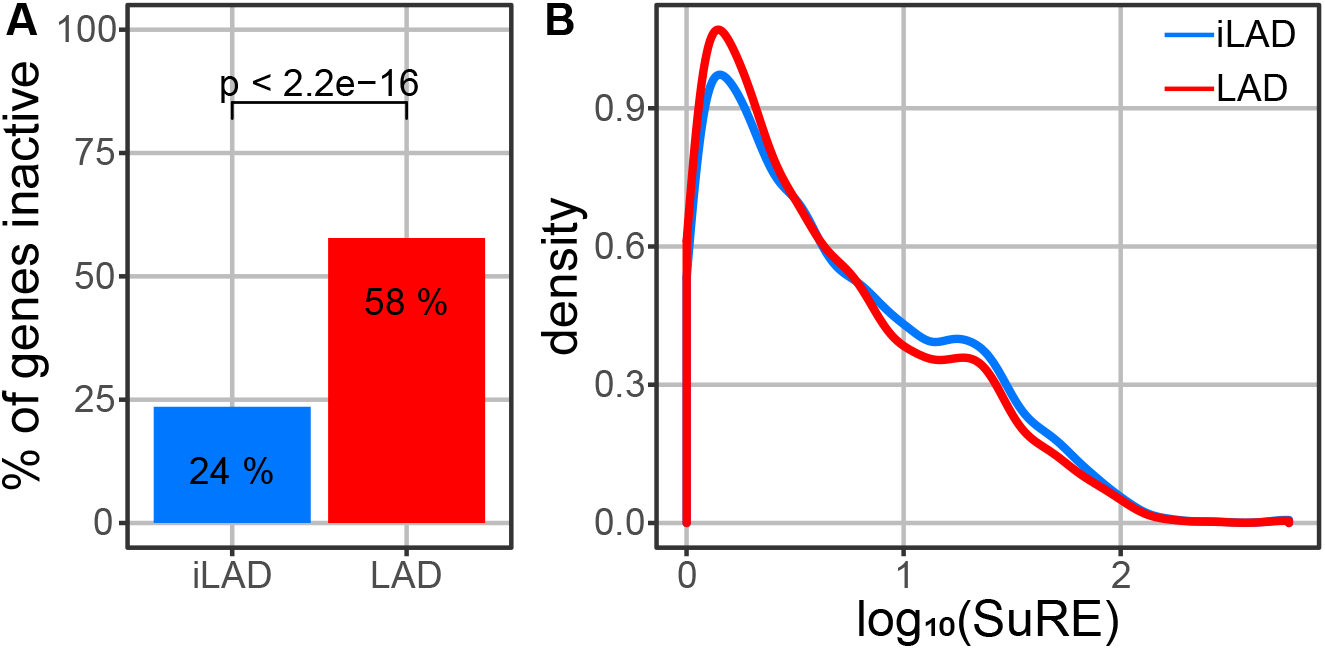
Promoters with SuRE activity inactive in endogenous context. **(A)** Percentage of promoters active in episomal plasmid context (log_10_(SuRE) > 0) with no measurable activity in native context (log_10_(GRO-cap) < -3) for LAD promoters and a subset of iLAD promoters matched on SuRE activity. **(B)** Density of expression in episomal plasmid context (log_10_(SuRE)) for matched promoter sets used for (A).

**Supplementary figure S2.**
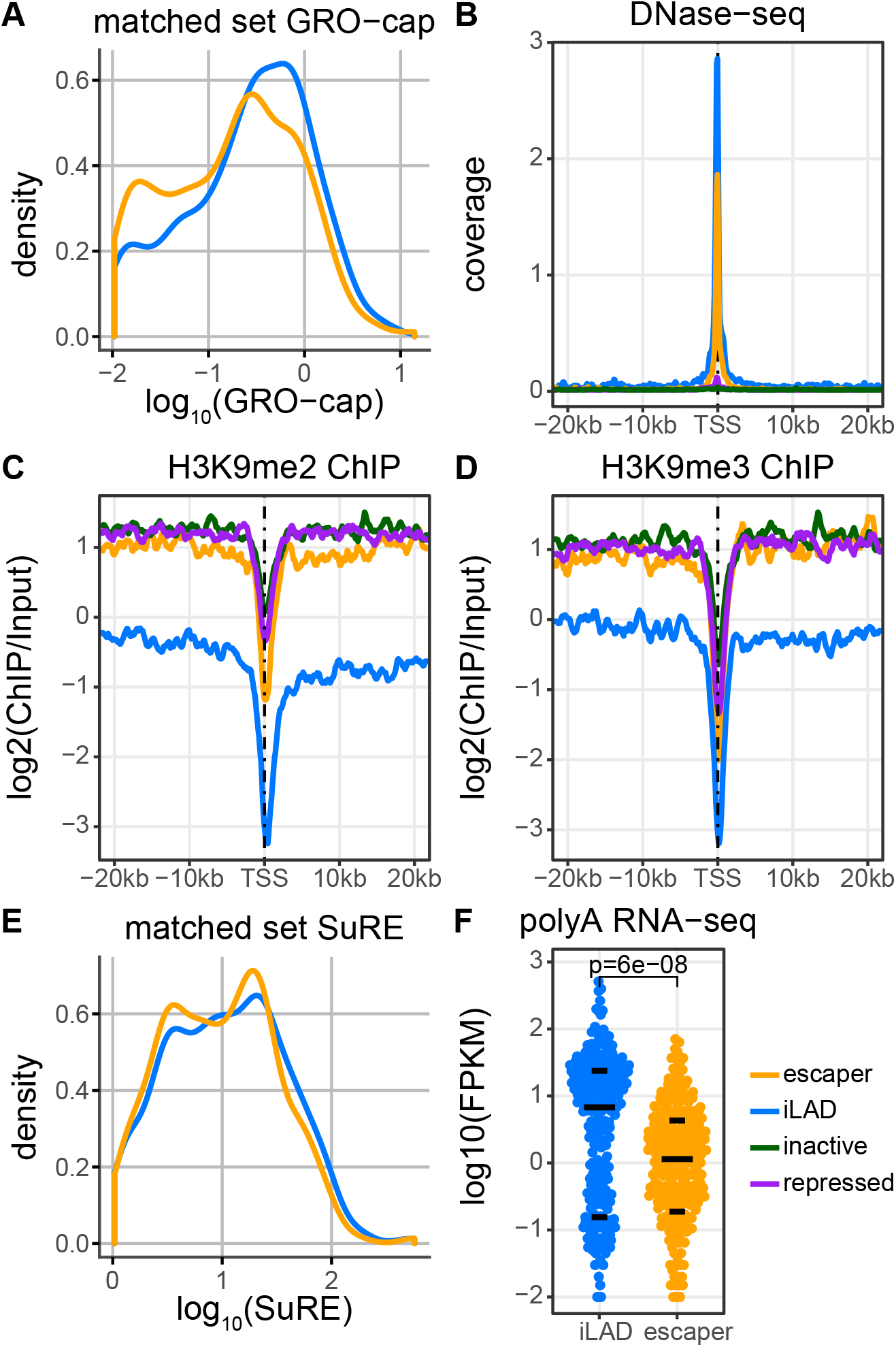
Properties of escaper promoters and their genes in native context. **(A)** Matched set of iLAD and escaper promoters used in Figure 2 and S2B-D. Promoters in iLADs were matched on GRO-cap activity of escaper promoters. **(B)** Average DNase-seq profile of each promoter class using paired-end sequencing. Data are from (Dunham et al., 2012). **(C)** Average H3K9me2 signal around each promoter class according to ChIP-seq (Salzberg et al., 2017). **(D)** Average H3K9me3 signal around each promoter class according to ChIP-seq (Salzberg et al., 2017). **(E)** Matched set of iLAD and escaper promoters used in Figure S2F. Promoters in iLADs were matched on SuRE activity of escaper promoters. **(F)** Same as Figure 2A, but with iLAD promoters matched on SuRE activity.

**Supplementary figure S3.**
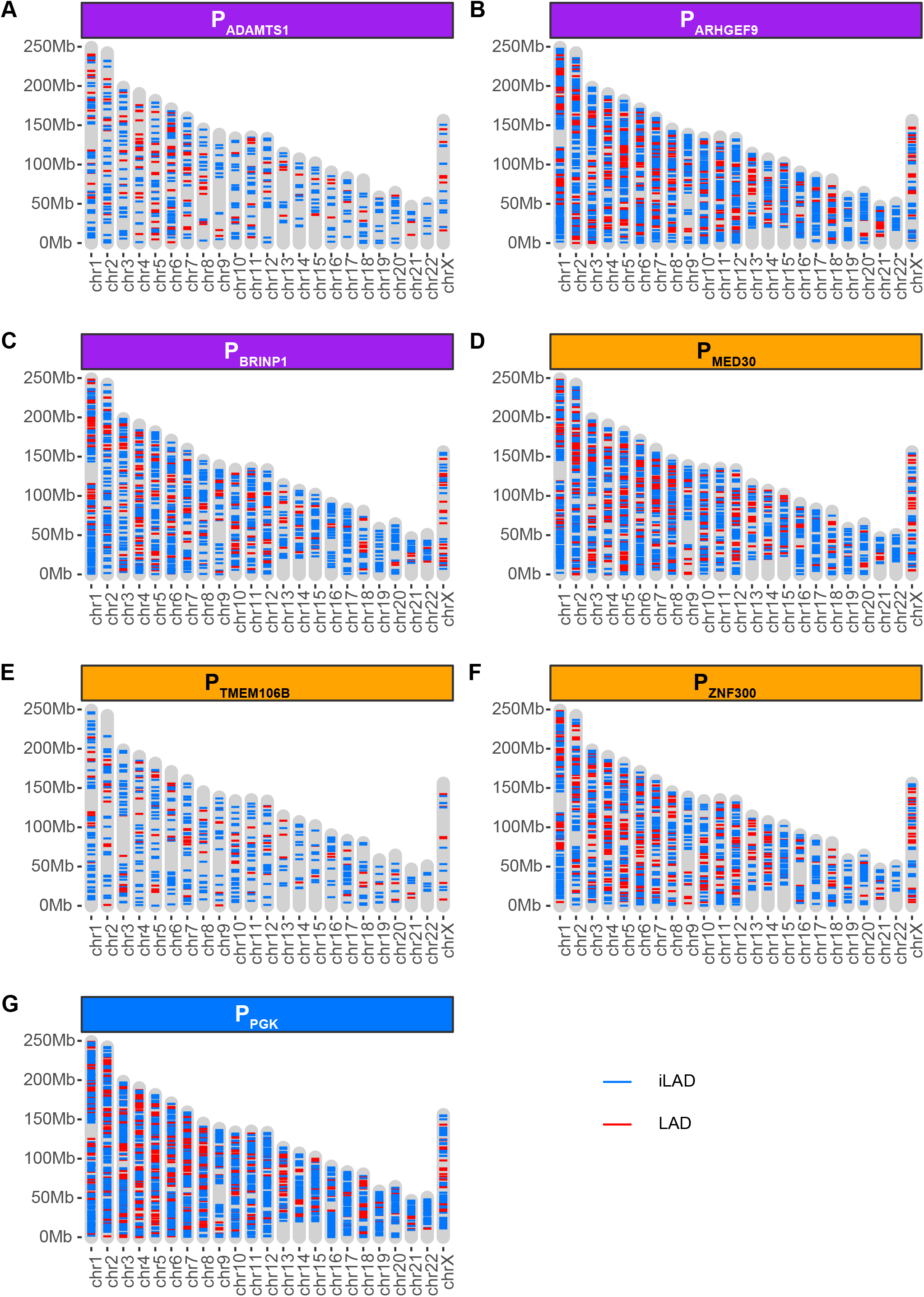
Chromosomal locations of TRIP integrations. **(A-G)** Locations of TRIP reporter integrations that could be uniquely mapped to the genome and assigned to a unique barcode, for each promoter. Red: integrations in LAD, blue: integrations in iLADs.

**Supplementary figure S4.**
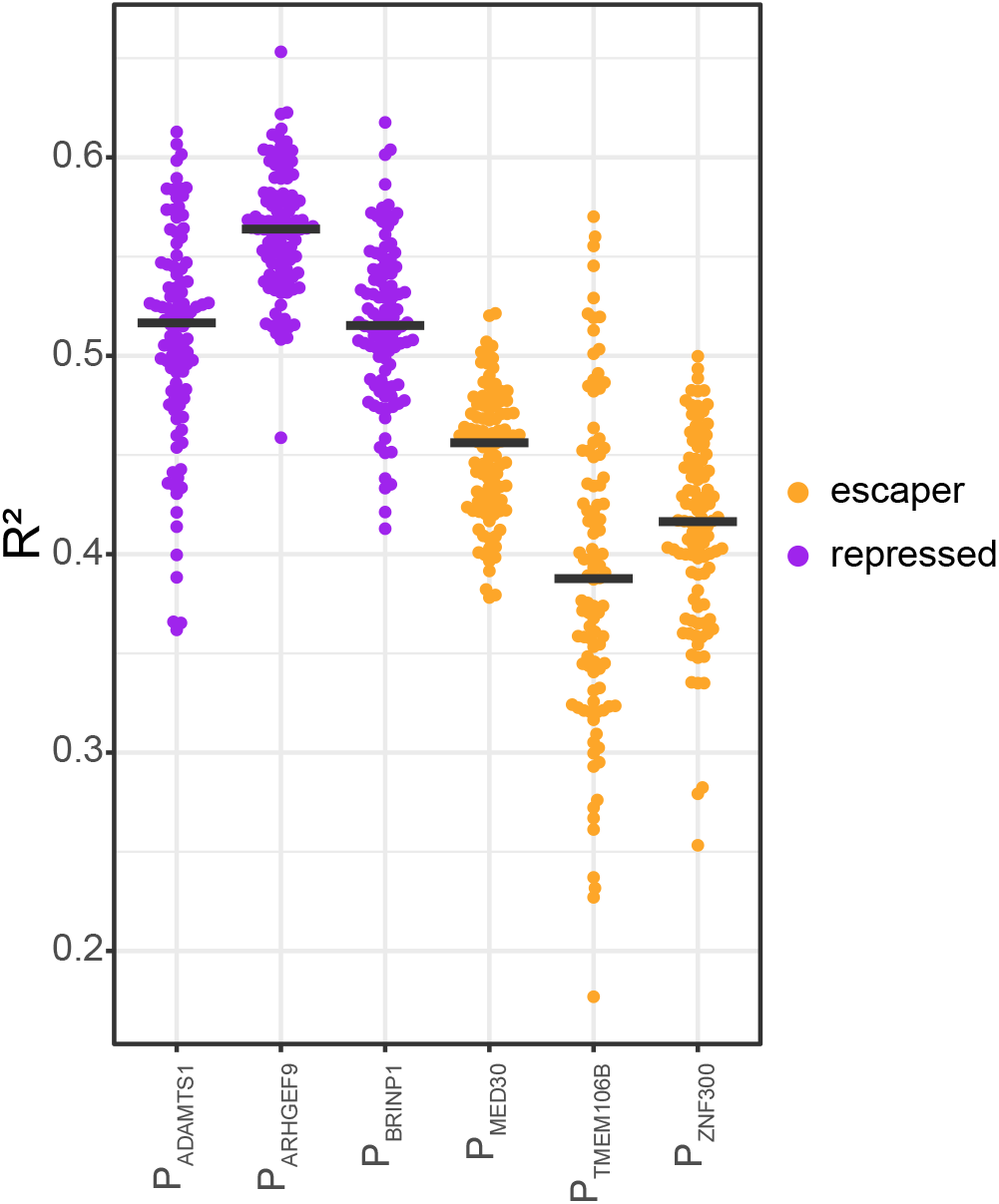
Modeling TRIP expression levels of individual promoters as function of chromatin features in all locations. R^2^ values of 100 lasso regression models of TRIP data for individual promoters using all available integrations (both LAD and iLAD). Each dot represents a model. Black lines represent the median R^2^ values.

## SUPPLEMENTARY DATASETS

Supplementary datasets were deposited on OSF: https://osf.io/myu3k/.

**Supplementary Dataset S1. SuRE activity of annotated promoters.**

Data are from (van Arensbergen et al., 2016) and were re-mapped to genome build hg38.

**Supplementary Dataset S2. TRIP activity of seven promoters at uniquely mapped integration sites.**

**Supplementary Dataset S3. SuRE activity of annotated enhancers.**

Data are from (van Arensbergen et al., 2016) and were re-mapped to genome build hg38.

## METHODS

### Promoter and enhancer definitions

For the comparison of GRO-cap and SuRE, promoters were taken from GENCODE (Harrow et al., 2012) version 27, with the following additional requirements. First, to focus on annotated promoters with likely functional relevance, we required that the promoters are active in at least one cell type or tissue according to the FANTOM5 database, reprocessed for hg38, using CAGE peaks lifted over from hg19 passing QC filters and CAGE peaks newly identified in hg38 (fair + new set), version 4 (Abugessaisa et al., 2017). For this criterion, only promoters with at least one CAGE peak within 50 bp were used. Second, to avoid overlapping data due to the resolution of SuRE, promoters were required to be at least 500 bp apart. For alternative promoters from the same gene located within 500 bp from each other, we selected the most active promoter, using the sum of the expression levels normalized by read depth for each experiment in the FANTOM5 database.

Enhancer coordinates were taken from the genehancer v4.8 database (downloaded from https://genecards.weizmann.ac.il/geneloc/index.shtml). We defined enhancer regions as 300 bp windows centered around the centre of these enhancers. We only used enhancers that were at least 5 kb away from any promoter as listed by Gencode v27.

### GRO-cap and SuRE

GRO-cap data (Core et al., 2014) was aligned to the hg38 genome using Bowtie2. Similar to (Core et al., 2014), reads were trimmed to 30 bases and first mapped to the human ribosomal DNA complete repeat unit (Genbank ID: U13369.1). Unaligned reads were then aligned to hg38 (without alternative haplotypes). Reads were aligned using Bowtie2. The *bamcoverage* tool from the *deeptools* package was subsequently used to create separate coverage tracks for forward and reverse strands. For this, using an offset and binsize of 1.

For SuRE data, reads were aligned to hg38 and processed as previously reported (van Arensbergen et al., 2016), but re-mapped to the hg38 version of the human genome. Average SuRE activity was computed within a 1kb window centered on each TSS or enhancer, similar to (van Arensbergen et al., 2016). A pseudocount equal to half the minimum expression was added before log_10_ transformation.

### LAD promoter and enhancer classifications

The three classes of LAD promoters were defined based on a combination of GRO-cap signals, SuRE signals, and a "LAD Repression Score" (LRS). The LRS is the deviation of the measured GRO-cap signal of promoters in LADs from the average GRO-cap signal of promoters in iLADs with a matching SuRE signal. This was calculated as follows. First, all promoters were sorted based on their SuRE expression. Next, for 60 windows of 501 promoters, equally distributed across the sorted list, the average log_10_(GRO-cap) and log_10_(SuRE) values for the iLAD promoters were calculated. A complete curve was then obtained by linear interpolation of the 60 points (blue curve in **Figure 1A,B**). Subsequently, for all LAD promoters the LRS was calculated by substracting the predicted GRO-cap score according to this curve from the measured GRO-cap score. Finally, LAD promoters with a measured log_10_(SuRE) value < -0.3 and a log_10_(GRO-cap) value < -2 were classified as *inactive*; LAD promoters with a log_10_(SuRE) value > 0.3, log_10_(GRO-cap) < -2 and LRS < -1 were classified as *repressed*. LAD promoters with a log_10_(GRO-cap) value > -2, a log_10_(SuRE) value > 0 and LRS > -0.5 were classified as *escaper*. The cutoff values are shown as dotted lines in **Figure 1C**.

A similar strategy was used to define the three classes of enhancers. Specifically, LAD enhancers with a measured log_10_(SuRE) value < -0 and a log_10_(GRO-cap) value < -2.8 were classified as *inactive*; LAD enhancers with a log_10_(SuRE) value > 0, log_10_(GRO-cap) < -2.8 were classified as *repressed*; LAD promoters with a log_10_(GRO-cap) value > -2 and a log_10_(SuRE) value > 0 were classified as *escaper*.

### Selection of promoters for TRIP

The *escaper* promoters for TRIP were selected to have at least log_10_(SuRE) > 0.5 and log_10_(GROcap) > -0.5 and to show clearly detectable expression in mRNA-seq data from K562 cells (ENCODE accession ENCSR000CPH (Dunham et al., 2012)). We excluded promoters with other nearby known active elements such as other promoters. Similar criteria were used to select *repressed* promoters except that these promoters were selected for low GRO-cap and mRNA-seq levels. Detailed information on the selected promoters is provided in **Supplementary Table S1**.

### TRIP plasmid construction and generation of TRIP cell pools

TRIP was essentially performed as described (Akhtar et al., 2014). The piggyBac reporter construct of (Akhtar et al., 2013) was used, except that the 14 GAL4 repeats, the mouse phosphoglycerate kinase (mPGK) promoter, the Puromycin resistance cassette (PuroR) and the internal ribosome entry site (IRES) were replaced by the promoter of interest. For each promoter, libraries of plasmids with 16 bp random barcodes were generated by electroporation into CloneCatcher DH5G electrocompetent *Escherichia coli* (#C810111; Genlantis) (*PGK*, *ADAMTS1*, *ARHGEF9*, *THEM106B*) or into E. cloni 10G (#60107-1; Lucigen) (*MED30*, *ZNF300* and *BRINP1*). Plasmid libraries had a complexity in the range of 40000 – 352500 (estimated by colony counting).

Next, 15 μg of plasmid DNA from each library was co-transfected with 5μg mCherry-expressing plasmid in 4 million K562 cells using Lipofection with Lipofectamine 2000 (ThermoFisher #11668019). Successfully transfected cells were obtained by FACS sorting of mCherry-positive cells, and cell pools carrying random reporter integrations were obtained as described (Akhtar et al., 2014).

### Analysis of TRIP expression data

For each promoter, two independent TRIP experiments were done with two independent cell pools each. Quantification of barcodes in cDNA and genomic DNA was performed as described (Akhtar et al., 2014), using Illumina HiSeq2500 sequencing with read length 75 bp.

Barcodes were extracted from cDNA and gDNA reads using an in-house script using functions from *cutadapt*. After demultiplexing, the start of the sequence was matched to the expected sequence GTCACAAGGGCCGGCCACAACTCGAG, allowing for an error-rate of less than 0.1 (1 mismatch per 10 bp). If this sequence matched the read, the subsequent 16 basepairs were classified as barcode and stored only if the sequence after this barcode matched the expected sequence TGATCCTGCAGTG with the same maximum error-rate as the preceding sequence. IUPAC codes in sequences are also considered as matching (e.g. N matches any nucleotide).

To identify the total set of genuine barcodes in each cell pool we used the barcodes extracted from the gDNA reads. We required each barcode to be present in at least 5 reads. To eliminate aberrant barcodes arising from mutations during PCR and sequencing, we applied *starcode* (Zorita et al., 2015), using sphere clustering with a maximum Levenshtein distance of 2.

Next, for the resulting set of genuine barcodes, the counts in both cDNA and gDNA reads were determined. A pseudocount of 1 was added to the cDNA counts. Subsequently, counts for both cDNA and gDNA were normalized to to the total respective barcode counts, and resulting cDNA values were divided by gDNA values. After log2-transformation these ratios were averaged across the two replicate experiments to obtain the expression score for each barcode. Data from two independent cell pools (each with different sets of integrations) were then combined.

### Mapping of TRIP reporter integrations

Mapping of genomic integration sites together with barcode identities was done by inverse PCR (iPCR) followed by 2 x 75 bp paired-end sequencing on an Illumina HiSeq2500 as described (Akhtar et al., 2014). Parsing of the reads was done as follows. The first read in each pair was used to extract the barcode. This was done by first removing GTCACAAGGGCCGGCCACAAC constant sequence and subsequently matching a regular expression TCGAG[ACGT]{16}TGATC. From this sequence, the 16 bp barcode was extracted. Barcodes were matched to the genuine set of barcodes obtained from gDNA sequencing reads and variations upon this set of barcodes likely caused by errors were discarded.

The second read of each pair was used to locate the site of integration after removing the GTACGTCACAATATGATTATCTTTCTAGGGTTAA region which matches the transposon arm. The flanking sequence was aligned to hg38 using *bowtie2* using the *very-sensitive-local* option (20 seed extension attempts, up to 3 re-seed attempts for repetitive seeds, 0 mismatches per seed, with a seed-length of 20 and using a multi-seed function *f(x) = 1 + 0.5 * sqrt(x)*). Locations of integration sites were required to be supported by at least 5 reads with an average mapping quality larger than 10 at the primary location, having at least 70% of the reads located at this locus, with not more than 10% of the reads at a secondary location.

### Multiplexed measurement of promoter activity in plasmid context

For each promoter, the barcoded TRIP plasmid library was re-transformed into DH5-alpha *E. coli* and plated; 100 colonies were picked, pooled, and expanded to produce plasmid mini-libraries with 100 different barcodes each. The barcodes in these mini-libraries were PCR amplified and sequenced by Illumina MiSeq to identify the barcodes belonging to each promoter. Next, the plasmid mini-libraries of the 7 promoters were mixed in equal porportions and transfected into K562 cells by the same nucleofection method as used for SuRE (van Arensbergen et al., 2016). Two days after transfection, barcodes were counted in cDNA by PCR amplification and Illumina sequencing, similar to TRIP. The same was done for the input plasmid DNA. For each barcode, cDNA counts were then normalized to plasmid counts to obtain the relative expression. The resulting values were log2-transformed, and results from two independent replicate experiments were averaged.

### LMNB1 DamID-seq

K562 cells were grown according to the ATCC culture protocol (https://www.lgcstandards-atcc.org). Cells were lentiviral transduced with Dam or Dam-LMNB1 constructs, both fused to a destabilization domain and under control of the PGK1 promoter (Amendola et al., 2005; Kind et al., 2013). A GFP sample was added to estimate transduction efficiency. To keep expression low, Dam fusion proteins were not stabilized with Shield1. Three days after transduction, gDNA was isolated using the Bioline Isolate II genomic DNA kit (BIO-52067). Additional RNAse A (4 uL of 10 mg/uL) was added during cell lysis.

DamID was performed similar to Vogel et al., 2007, but modified for next generation sequencing as described below. 500 ng of DNA was digested with DpnI (0.5 uL DpnI (20U/uL New England Biolabs #R0176L), 1 μl 10x CutSmart) in 10 uL total volume for 8 hours at 370C, followed by 20 minutes of heat inactivation at 800C. Adapters were ligated to the digested fragments by adding 10uL of ligation mixture (0.5 uL T4 ligase (5U/μl Roche #10909246103), 2 uL 10x ligation buffer, 0.25 uL DamID adapter (dsAdR 50 μMVogel et al., 2007) and 7.25 uL H2O). After an overnight incubation at 16C, ligase was heat inactivated for 10 minutes at 650C. Unmethylated GATC sequences were cleaved with DpnII (addition of 1 μl DpnII (10U/μl New England Biolabs R0543L), 5 μl DpnII buffer and 24 μl H2O) by incubating 1 hour at 370C. During these steps, controls without DpnI and ligase were included to assess specificity of the amplification. 8 uL of the final digestion was added to a PCR mixture (20μl 2xMyTaq (Bioline BIO-25041), 1 uL primer (Adr-PCR-Rand1, 50 μM) and 11 uL H2O) and amplified with the following PCR protocol: 8 minutes at 720C, cycles of 20 seconds 940C, 30 seconds 580C and 20 seconds 720C, and finally 2 minutes at 720C. 23 and 22 amplification cycles were used for replicates 1 and 2, respectively. 4 uL of the PCR mixture was put on gel to verify specific amplification, while the rest was column purified using Bioline Isolate II PCR and Gel kit (Bioline BIO-52060).

Sequencing libraries were prepared by a series of end-repair in 50 uL (End-It, Lucigen, ER81050), column purification (Bioline BIO-52060), Klenov 3A-overhang in 50 uL (Klenov 3’ -> 5’ exo-, New England Biolabs, M02125) and bead purification (1.8x CleanPCR beads, CleanNA, CPCR-0050), all following provider’s protocols. DNA concentration was measured by Nanodrop and ~250 ng of DNA was resolved in 6.5 uL H2O. Y-shaped adapters were ligated overnight at 160C, by adding 3.5 uL ligation mixture (0.5 μl T4DNA ligase (5U/μl Roche #10909246103), 1 μl 10x ligation buffer, 0.5 μl Y-adapter (50 μM) and 1.5 uL H2O), followed by 10 minutes heat inactivation at 650C and another round of bead purification resolving in 20 uL H2O (1.8x CleanPCR beads, CleanNA, CPCR-0050). Illumina indices were added with amplication, using 8 uL bead purified DNA in a 20 uL PCR reaction (10 uL 2xMyTaq (Bioline BIO-25041), 0.5 uL Illumina P5 primer (5.0 μM), 0.5 uL Illumina P7 indexing primer (5.0 μM) and 1 uL H2O). The following PCR protocol was used: 1 minute 940C, cycles of 30 seconds 940C, 30 seconds 580C and 30 seconds on 720C, followed by 2 minutes at 720C. 10 and 11 cycles were used for replicates 1 and 2, respectively, and 4 uL was put on gel. Based on smear intensities, samples were pooled and cleaned with bead purification (1.6x CleanPCR beads, CleanNA, CPCR-0050). Samples were sequenced on a HiSeq 2500 with High Output Mode of single-end 65 bp reads.

### DamID-seq data processing

First, the constant sequence of the DamID adapter was trimmed from the 65 bp single-end reads (with cutadapt 1.11 and custom scripts). The remaining gDNA starting with GATC was mapped to a combination of GRCh38 v15 (without alternative haplotypes) and a ribosomal model (GenBank: U13369.1) with bwa mem 0.7.17. Reads with a mapping quality of at least 10 were counted to GATC fragments. The middle of the GATC fragment was used to combine these counts into bins of various sizes. Only bins with at least 10 reads (combined target + Dam-only) were subsequently normalized. These bins were first normalized to 1M reads and with a pseudocount of 1 a log2-ratio over the Dam-only control was calculated. LADs were defined by running a hidden markov model over the normalized values (using the R-package HMMt; https://github.com/gui11aume/HMMt). BigWigs tracks were generated of the normalized counts per GATC-fragment. These normalized BigWigs were used to calculate average count in 100bp bins around the TSS using the computeMatrix function from deeptools. These bins were used to calculate a running sum using the sum of 5 bins for each 100bp bin. These running sums were then used to calculate a fold-change of the LmnB1-Dam over Dam signal.

### Chromatin immunoprecipitation data analysis

We re-processed published ChIP-seq data from various sources (**Supplementary Table S3**) for consistency. Raw sequencing data was obtained from the sequence read archive (SRA). Reads were aligned to the human genome hg38 using *bowtie2* with default options. Replicates for sample data were processed separately, while the sequences from the input were combined. After alignment, reads were filtered on a minimum mapping quality of 30 and duplicate reads were removed except for the reads coming from experiments using tagmentation, in which duplicates were kept. After this, regions were masked based on blacklist regions identified by the ENCODE project (ENCFF419RSJ) (Dunham et al., 2012) and artifact regions identified based on the input reads using chipseq-greylist, a python implementation of GreyListChIPs (Gordon Brown, 2018). We considered ChIP-seq datasets to be of sufficient quality for our analyses if there was well annotated input and sample data available and consistent read lengths were used. Filtered alignments were used to call domains with significant enrichment by hiddenDomains with a binsize of 1kb and minimum posterior of 0.9. Overlapping domains were selected between replicate experiments. For the calling of endogenous genes located in H3K27me3 regions, binsize was set to 20kb and a posterior of 0 was used.

Mean Chip-seq signals for specific regions (e.g. TSSs, genebodies, TRIP integration sites) were calculated by taking the sum of the reads in the region, scaling input and sample counts by the smallest library size, adding a pseudo count of 1 and subsequently dividing sample over input normalized counts. After this, replicate experiments were averaged. Enrichment signal along a window around sites of interest (e.g. TSS) was calculated by first using *computeMatrix* function of the *deeptools* package to calculate read coverage in 100bp bins centered around the site of interest(Ramírez et al., 2016). After calculating coverage, smoothed average signal for each group of interest was calculated by using running mean with a width of 9 bins. Coverage was then normalized by library size and enrichment over input was calculated.

### Calling domains of TRIP integrations

TRIP integration sites were called as either LAD or H3K27me3 localized when the TTAA sequence of integration falls within either domain. Overlap was calculated using the *intersect* function of *bedtools* (Quinlan and Hall, 2010). For H3K27me3 domains both datasets by (Schmidl et al., 2015) were used and integrations overlapping with either dataset were classified as H3K27me3 domain integrations.

### Calling of endogenous domains

Endogenous genes were called as either LAD or H3K27me3 localized when the complete gene-body fall within respective domains and are at least 1 binsize (5kb and 20kb respectively) from the border. Overlap was calculated using the *intersect* function of *bedtools* (Quinlan and Hall, 2010). For H3K27me3 domains, unlike previous TRIP data, overlap was calculated based on individual replicates, genes completely overlapping with at least 2 replicates were called as located inside H3K27me3 domain.

### TT-seq

TT-seq data from K562 cells (Schwalb et al., 2016) was kindly provided by B. Schwalb and P. Cramer as 4 BigWig files, two for each replicate (one file for forward and one for for reverse orientation relative to the hg38 reference genome). Windows around the TSS were created similarly to the approach used for the DamID-seq and ChIP-seq data by first using the *computeMatrix* function. For each TSS, the bins in sense orientation with its gene were used to calculate running means with a width of 3 bins. Subsequently the average for each class and across 2 replicates was calculated.

### Statistical modeling of TRIP and epigenome data

For statistical modeling of TRIP expression, mean ChIP-seq and DamID signals were calculated within a region centered on the integration site (5kb and 10kb respectively). Mean values were averaged between experiments. Proximity to the nearest domain was calculated as –*log*_10_ (*distance*+1). For experiments with the same target, the minimum distance was used. For integrations inside LADs, the proximity to nearest border was calculated.

To predict barcode expression values (log2), L1-regularized linear regression models (lasso) were trained using mean ChIP-seq/DamID signals at the integration sites along with proximity features. Only TRIP integrations within LADs were considered for each promoter class (escaper or repressed). A total of 100 models were fitted for each class. For each model, data points were first randomly split into a training set (80%) and a test set (20%). The model was trained with 10-fold cross-validation using the glmnet package v2.0-16. The value of the regularization parameter that minimized the cross-validated mean squared error was used for model selection. The prediction accuracy (R^2) of the selected model was computed on the test set. For promoter-specific models, both LAD and iLAD integrations were considered.

Feature importance analysis was performed using bootstrap-Lasso essentially as described in (Comoglio and Paro, 2014). The importance of a feature corresponds to its selection probability (normalized frequency of non-zero coefficients) across all bootstrap-Lasso models. All features with a selection probability > 0.7 were considered.

### Sequence motif analysis

For the motif analysis, enhancers identified as escapers were compared to a subset of enhancers of the repressed class, matched on SuRE activity. All escaper and repressed enhancers were sorted on SuRE expression and subsequently all enhancers ranking right before and after each escaper enhancer in the list were selected. Of this group of enhancers, the repressed enhancers were used for the matching set.

*DREME* was used to discover denovo motifs enriched in escapers with repressed enhancers as background(Bailey, 2011). The number of most significant words passed in the beam search was set to 1000 and the maximum word-size was set to 10. The other settings were left to default (E-value Threshold 0.05, minimum word-size of 3). Both forward and reverse strand were processed.

For each enhancer the 300bp region around the centre was extracted from the human genome version hg38 using the *getfasta* command from *bedtools* (Quinlan and Hall, 2010). Subsequently, for a well curated set of motifs, we approximated position specific affinity matrices (PSAM) by transforming the position weight matrix (PWM) by dividing each column, representing 1bp, by it’s maximum weight value. *AffinityProfile* from the *REDUCE Suite* was used to calculate the sum of affinities calculated along both orientations of the 300bp sequence(Roven and Bussemaker, 2003). Motifs were classified as either pioneer or non-pioneer based on the classification by Ehsani *et al.*(Ehsani et al., 2016). For every motif a median affinity was calculated for each enhancer class and the median for escapers was divided by the median for repressed. Only TFs that are detectably expressed in K562 cells (RNA FPKM > 1 in ENCODE mRNA-seq dataset ENCSR000CPH) were included.

---

**Supplementary Table S1.** Specifications of promoters used for TRIP experiments.

| **Gene Symbol** | **promoter region used** | **transcript ID** | **gene ID** | **class** |
| --- | --- | --- | --- | --- |
| **ADAMTS1** | chr21:26845360-26847082 | ENST00000284984.7 | ENSG00000154734.14 | repressed |
| **ARHGEF9** | chrX:63755062-63756421 | ENST00000624538.1 | ENSG00000131089.15 | repressed |
| **BRINP1** | chr9:119369375-119370353 | ENST00000373964.2 | ENSG00000078725.12 | repressed |
| **MED30** | chr8:117519681-117520764 | ENST00000297347.7 | ENSG00000164758.7 | escaper |
| **TMEM106B** | chr7:12210274-12211294 | ENST00000442107.1 | ENSG00000106460.18 | escaper |
| **ZNF300** | chr5:150904928-150906439 | ENST00000394226.2 | ENSG00000145908.12 | escaper |
| **PGK** | chrX:78103813-78104327 | ENST00000373316.4 | ENSG00000102144.13 | iLAD |

**Supplementary Table S2.**
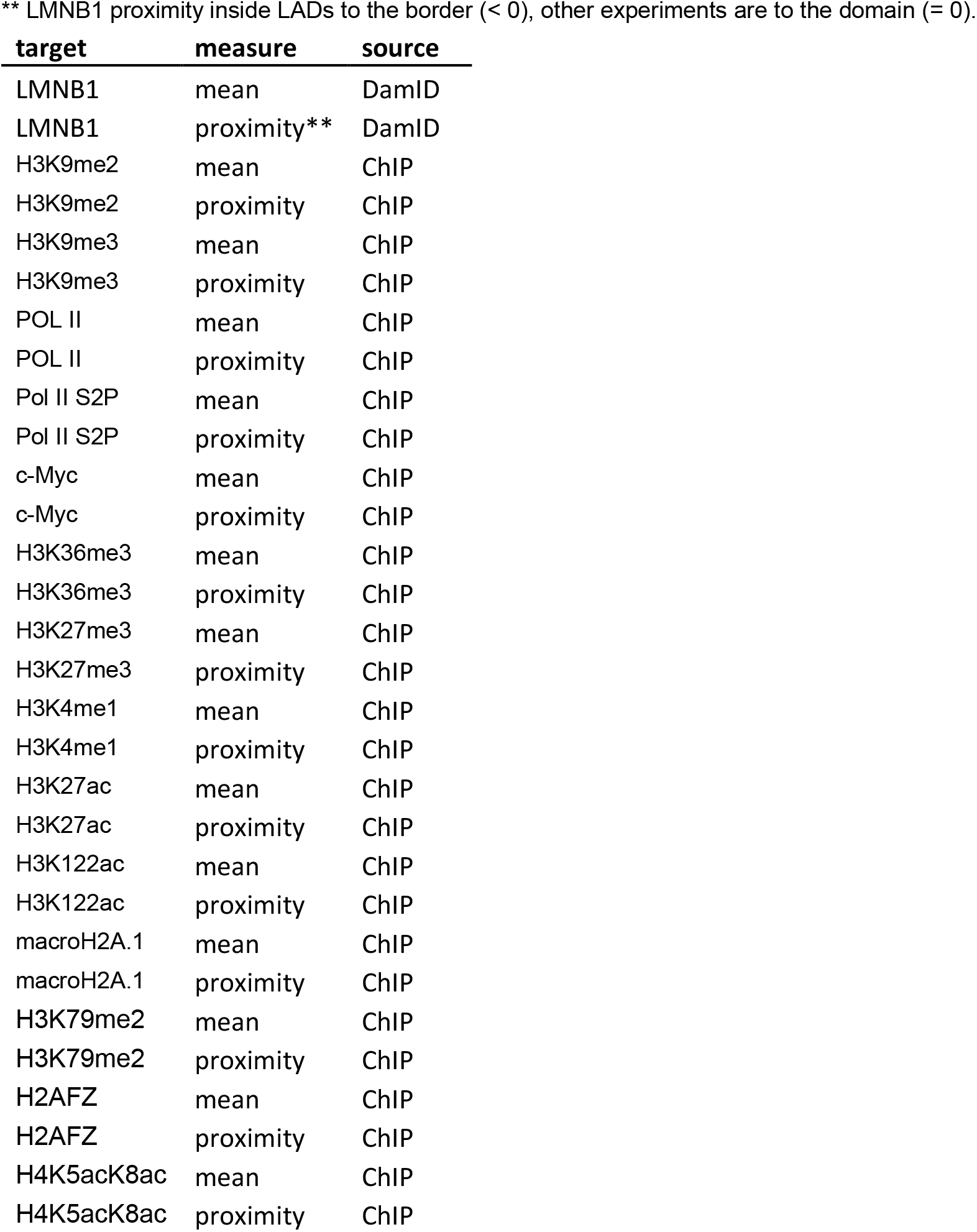
Chromatin features used as input for statistical modeling. ** LMNB1 proximity inside LADs to the border (< 0), other experiments are to the domain (= 0).

**Supplementary Table S3.**
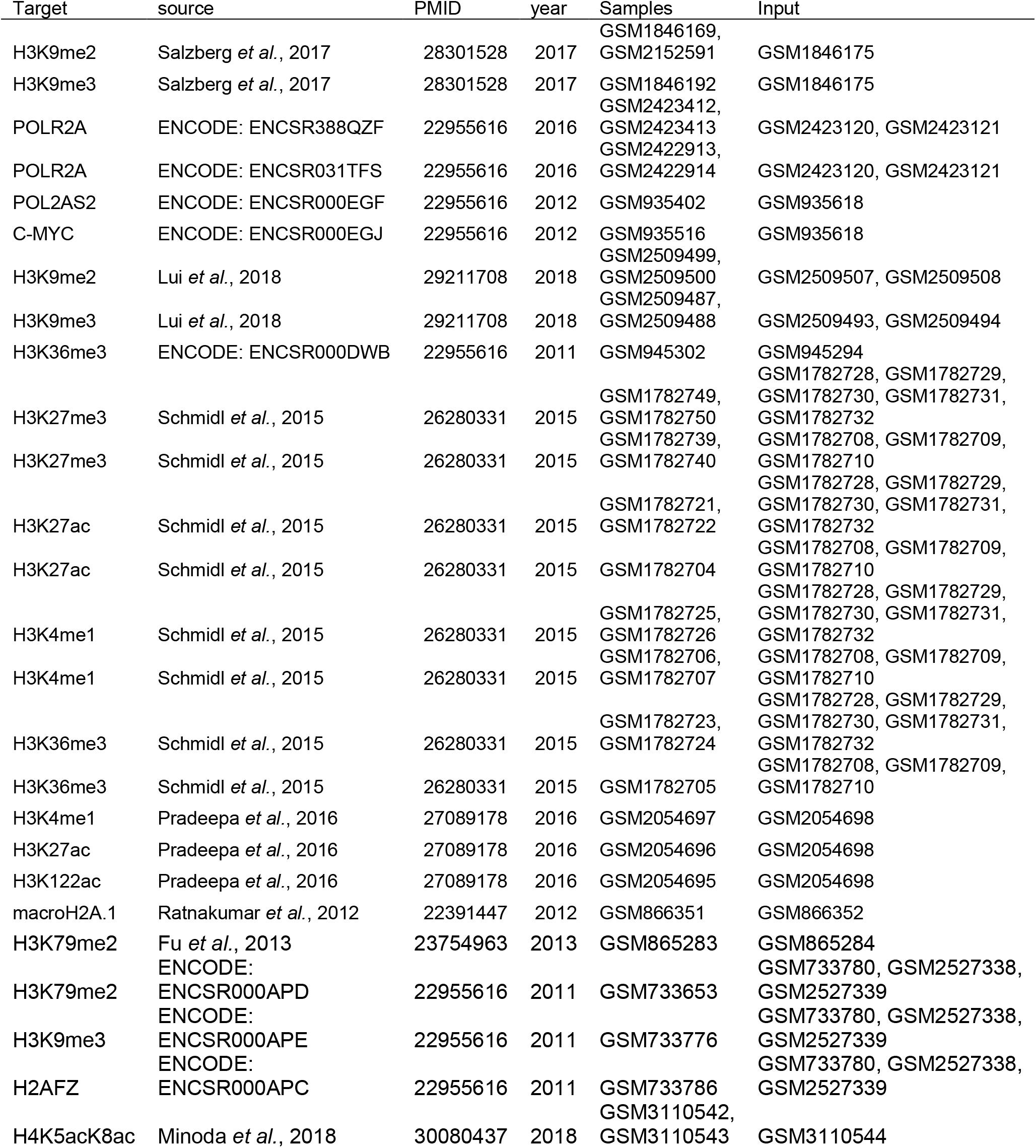
ChIP-seq datasets used.

**Supplementary Table S4.**
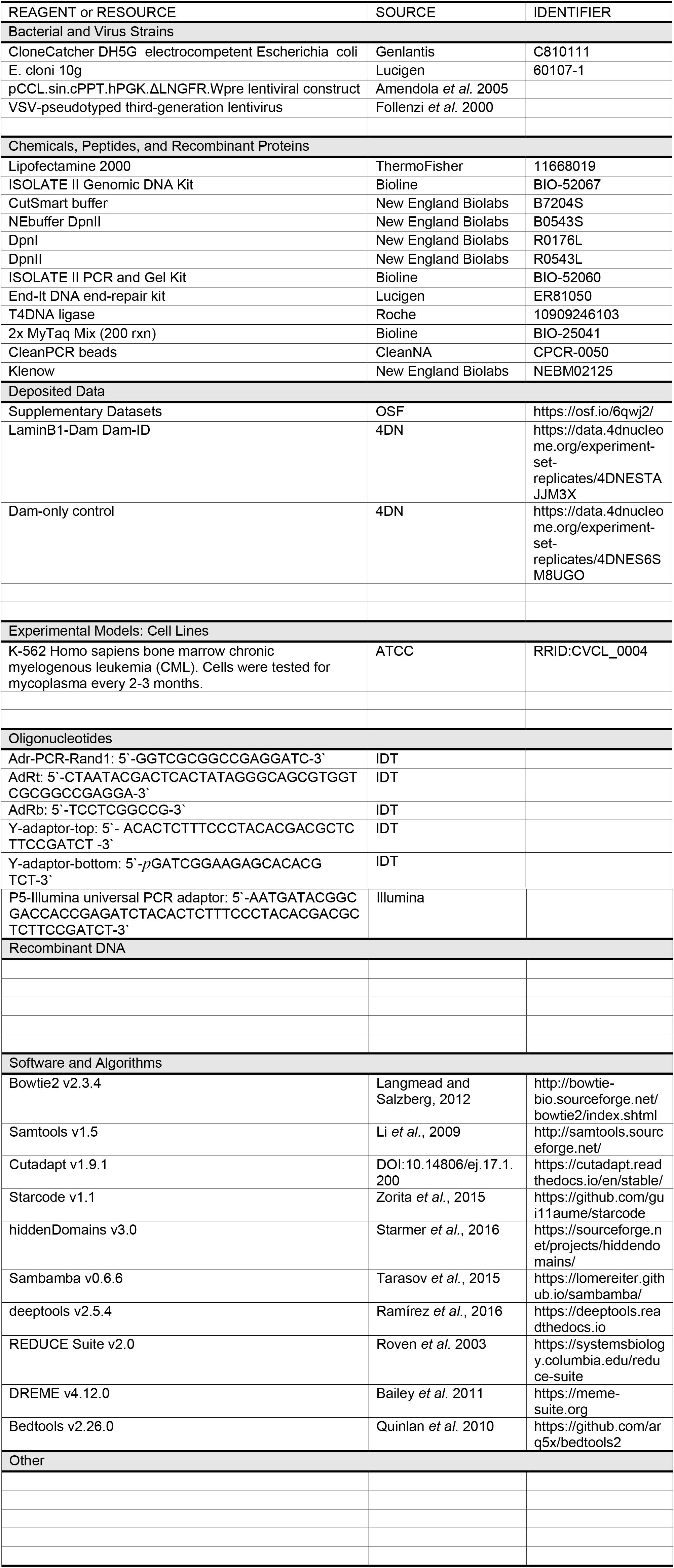
Reagents and software.

